# Novel Plasma Proteomics and Phosphoproteomics Platform Captures Pleiotropic Cardiometabolic Spectrum Effects of Semaglutide in Patients with T2D and Atherosclerosis: A Companion Diagnostic Pilot Study from the STOP (Semaglutide Treatment On coronary atherosclerosis Progression) Randomized Trial

**DOI:** 10.64898/2026.02.16.706239

**Authors:** Antigoni Manousopoulou, Cory H. White, Sajad Hamal, Raj Nihalani, Matthew J. Budoff, Spiros D. Garbis

**Author notes:** Co-first, equal contribution (Antigoni Manousopoulou, Cory H. White).

## Abstract

**BACKGROUND:** As a GLP1 R agonist, semaglutide is known to exhibits pleiotropic health effects across the cardiometabolic spectrum in patients with type 2 Diabetes Mellitus (T2D). However, in depth and unbiased protein and phosphoprotein level evidence that reflects such effects of semaglutide in plasma remains elusive.

**OBJECTIVES:** This pilot study applied an innovative plasma proteomics and phosphoproteomics technology to a sub-set of patients with T2D that participated in the Semaglutide Treatment effect on coronary atherosclerosis Progression (STOP) randomized trial. The aim of this study was to identify the systemic effects of semaglutide treatment in pathways that underpin its pleiotropic cardiometabolic spectrum health benefits.

**METHODS:** The study applied a proprietary liquid biopsy discovery proteomics platform and its derivative cardiometabolic spectrum database (International patent PCT/US2021/063407) to 16 patients from the STOP randomized trial. Plasma samples from 8 patients in the active group and 8 patients in the placebo group at baseline and 52 weeks post treatment were analyzed. The methodology entailed the use of a unique liquid fixative chemistry to instantly solubilize and stabilize plasma proteins and phosphoproteins at room temperature followed by their direct microflow monolithic partition chromatography, dialysis purification, solution phase proteolysis, multiplex isobaric stable isotope labeling of proteotypic peptides, lab-on-chip TiO2/ZrO2 phosphopeptide enrichment and nanotechnology enhanced ultra-high resolution LC-MS analysis. To identify differentially abundant proteins (DAPs) and phosphoproteins (DApPs) in patients treated with semaglutide vs. placebo, the respective abundance ratio for each was considered repectively. Ratios were log2-transformed to normalize their distribution. A one-sample T-Test (paired) using the two-stage Benjamini Yekutieli Krieger step up method for multiple hypothesis testing FDR correction of the p-value was performed. The threshold of significance was set at q≦0.05. DAPs/DApPs were corrected for placebo. The expressed proteome and phosphoproteome were further interpreted with a multifactorial computational biology pipeline to deconvolute their underlying protein-level molecular pathways and their networks along with transcriptional factors and kinases that regulate them.

**RESULTS:** This study achieved an extremely high depth-and-breadth in quantitative proteome and phosphoproteome coverage from only 20µL whole plasma equivalent from each patient. Namely, a total of 13,173 proteins and 25,578 phosphopeptides were fully profiled (*q≦ 0.05*). Of these, 1,040 were differentially abundant proteins (DAPs) and 1,064 were differentially abundant phosphoproteins (DApPs), in the semaglutide treated group after correcting for placebo, at an absolute log_2_-fold-change of ≧ 0.5, CV ≦ 15%, q*≦ 0.05*. Of interest, this study profiled over 85% of all proteins/phosphoproteins (6,700) reported to date based on the use of the well curated and up-to-date PaxDB database. Over 70% of these known proteins were of exosomal origin. Importantly, an additional ∼9000 plasma proteins and phosphoproteins of this study constituted entirely novel observations. Contextualization of the DAPS, DApPs, kinases and transcription factors for all statistically significant enriched canonical pathways (FDR-corrected *q<0.001*) revealed a wide array of pathophysiological processes attributed to semaglutide treatment. Furthermore, these pathways provided a molecular understanding to the reported imaging biomarkers against the same STOP trial patients.

**CONCLUSIONS:** This feasibility study demonstrated how an effective plasma proteomics and phosphoproteomics platform can generate a treatment-adaptive companion diagnostic molecular signature that holistically captures the multiple cardiometabolic spectrum health effects of semaglutide in patients with atherosclerosis and T2D.

## INTRODUCTION

Glucagon-like peptide-1 receptor agonists (GLP-1RAs), including semaglutide, have markedly advanced the management of cardiometabolic disease, extending their therapeutic impact well beyond glycemic control in type 2 diabetes mellitus (T2D) and weight reduction in obesity (1,2). Accumulating evidence from clinical trials highlights the pleiotropic benefits of semaglutide across the cardiometabolic spectrum, with demonstrated improvements in renal outcomes (3), hepatic function (4), and cardiovascular health (5). Supported by cardiac computed tomography (CT), prior studies have identified imaging biomarkers indicating that semaglutide treatment in patients with T2D favorably influences cardiovascular structure and disease burden (6,7), as well as hepatic steatosis (8).

Of particular clinical interest, proteins—through both their total abundance and post-translational modifications such as phosphorylation—govern fundamental biochemical pathways and interconnected signaling networks in human physiology. Proteins further integrate genetic information with epigenetic regulatory mechanisms, including transcription factors and kinases, that are implicated in cardiovascular and other chronic disease states (9,10). Accordingly, discovery-based proteomics enables the unbiased interrogation of thousands of proteins in tissue and liquid biopsies (e.g., plasma or serum, ascitic fluid, cerebrospinal fluid), offering a systems-level approach to elucidate the molecular mechanisms underlying semaglutide-mediated cardiometabolic protection, as well as potential off-target effects.

As originally articulated by Claude Bernard, the *milieu intérieur*—exemplified by blood plasma—represents an ideal matrix for probing internal physiological regulation, homeostatic balance, and its perturbation by disease and therapeutic intervention (11). From this vantage point, plasma constitutes a dynamic circulating compartment that reflects secretory inputs from multiple organs and tissues under specific physiological or pathological states. This property is, in part, attributable to plasma exosomes, whose molecular cargo provides an integrated systems snapshot of the cells and tissues from which they originate. Notably, exosome biogenesis is increased under conditions of chronic low-grade inflammation and metabolic stress associated with obesity, diabetes, and cardiovascular disease (12,13). In these settings, exosomes exhibit pronounced disease-context specificity and function as both organotypic messengers and organotropic mediators, facilitating the transfer of inflammatory and metabolic signals between cells and tissues, thereby amplifying disease propagation (13–16). This paradigm reinforces the concept that organs do not operate in isolation but instead function synergistically in health and disease manifestation, a defining hallmark of the human physiome and pathophysiome.

To address the analytical challenges inherent in comprehensive plasma proteome and phosphoproteome profiling, the authors developed the Biofluid Total Analytic System (BioTAS), an internationally patented, affinity-independent liquid biopsy discovery proteomics and phosphoproteomics platform, together with its derivative cardiometabolic spectrum proteome database (17). BioTAS is uniquely suited to extract, solubilize, and enrich proteins and phosphoproteins contained within plasma exosomes without the need for labor-intensive, technically complex, and reproducibility-limited isolation procedures (18, 19). The methodological foundations of the BioTAS platform and its application to clinical observational and interventional studies have been previously reported (20–32). To help decipher resulting analytical outputs, the BioTAS platform also employs PROMINIA (PROtein MINing Integrated Algorithms), an advanced computational biology pipeline. In addition to statistical modeling, PROMINIA robustly identifies disease-adaptive differentially abundant proteins and phosphoproteins with their regulatory transcription factors, and kinases to finally deconvolute their associated signaling pathways (33–44). Publicly available clinical and pharmacologic datasets, together with in-house generated Proteas platform databases across multiple disease states, were employed as training sets for PROMINIA. Collectively, these attributes enable precise, sensitive, and comprehensive interrogation of the plasma proteome and phosphoproteome to advance mechanistic understanding of disease pathophysiology and therapeutic response.

In the present pilot study, we demonstrate the utility of the BioTAS platform to plasma samples obtained from the Semaglutide Treatment Effect on Coronary Atherosclerosis Progression (STOP) randomized trial (7). The primary objective of this analysis was to identify pathophysiologically relevant molecular pathways that complement the previously reported imaging biomarkers (6–8), thereby providing mechanistic insight into their observed morphological and compositional features. Furthermore, the identification of key hub proteins and phosphoproteins within these signaling networks offers the potential to elucidate the multifaceted cardiometabolic effects of semaglutide. Integration of advanced imaging and systems-level proteomics may ultimately facilitate the development of personalized therapeutic strategies. Ultimately, a validated plasma-based protein and phosphoprotein biomarker panel could support the development of a companion diagnostic assay tailored to monitor response to semaglutide therapy, assess equivalency in safety and efficacy across dosing regimens—including injectable versus oral formulations—and inform the selection of adjunctive lifestyle or pharmacologic interventions relevant to cardiometabolic disease.

This proof-of-concept investigation was designed to establish a systems-level understanding of the clinical effects of semaglutide while defining the mechanistic architecture underlying its surrogate cardiometabolic benefits, including reductions in plasma glucose and HbA1c levels, weight loss, and favorable imaging biomarkers. By integrating quantitative plasma proteomic and phosphoproteomic profiling with upstream regulatory networks encompassing transcription factors and kinases, we sought to delineate signaling pathways selectively modulated by semaglutide relative to placebo. This integrative approach enables identification of functionally relevant protein and phosphoprotein signatures that may serve as the basis for a companion diagnostic platform, providing a high-resolution molecular snapshot of how GLP-1 receptor agonists attenuate systemic and cardiovascular injury.

## MATERIAL AND METHODS

### Plasma procurement

The plasma samples were obtained from the clinical cohort examined in the STOP randomized trial (7). Specifically, plasma samples from 8 patients in the active group at baseline and 52 weeks post treatment and from 8 patients in the placebo group at baseline and 52 weeks post treatment were subjected to proteomic analysis. The clinical characteristics of the cohort at baseline are presented in Supplementary Tables 1-3. A volume of 100μL of whole plasma from each specimen was aliquoted and vortex mixed for 30 sec with 400μL of Proteas’ proprietary liquid fixative solution before analysis.

### Proteomic sample preparation

A total volume of 80μL from each liquid fixative treated plasma extract was subjected to automated serial fractionation for its protein and native peptide content using a variant of the open tubular chromatography array lab-on-chip originally reported by the authors (21) and most recently subject matter of Proteas international patent (17) and automation enhancements (22). This lab-on-chip fractionation device is uniquely amenable to work with the plasma liquid fixative chemistry. The eluents were ultimately dialysis exchanged to 0.5 M TEAB and 0.05% SDS, measured for its protein content and subjected to reduction, alkylation and LysC/Trypsin proteolysis at a ratio of 1:30. The resulting proteotypic peptides/phosphopeptides were labeled with the TMTpro isobaric stable isotope reagents, as specified by the manufacturer (Thermo Pierce, Rockford, IL). Additional details could be found in the Supplementary Methods.

### Phosphopeptide isolation and enrichment

The TMTpro labeled peptide/phosphopeptide samples was subjected to step-gradient chromatography with an open tubular Lab Chip surface functionalized with TiO2/ZrO2 chemistry as reported by the authors (21) with additional design improvements in Proteas’ international patent (17). Further details could be found in the Supplementary Methods.

### Mass Spectrometry Analysis

Samples reconstituted in LC buffer A (0.1% formic acid in water), randomized, and then injected onto the state-of-the-art EASY-nLC 1200 ultra-high-performance liquid chromatography coupled to an ultra-high resolution Exploris 480 quadrupole-Orbitrap mass spectrometer (Thermo Fisher Scientific) with internal calibration and retrofitted with nano-spray ionization source and the Field Asymmetric Ion Mobility Source (FAIMS). Its advanced analytical features are described in recent review (45). Peptides were separated by a custom reverse phase analytical column integrated with nanospray emitter (Mixed mode C4 and C18, 1.6 µm particles, 100 Å pore, 50 µm ID × 25 cm L). Additional details could be found in the Supplementary Methods. Figure 1 illustrates an overview of the proteomic platform applied for this pilot study.

**Figure 1.**
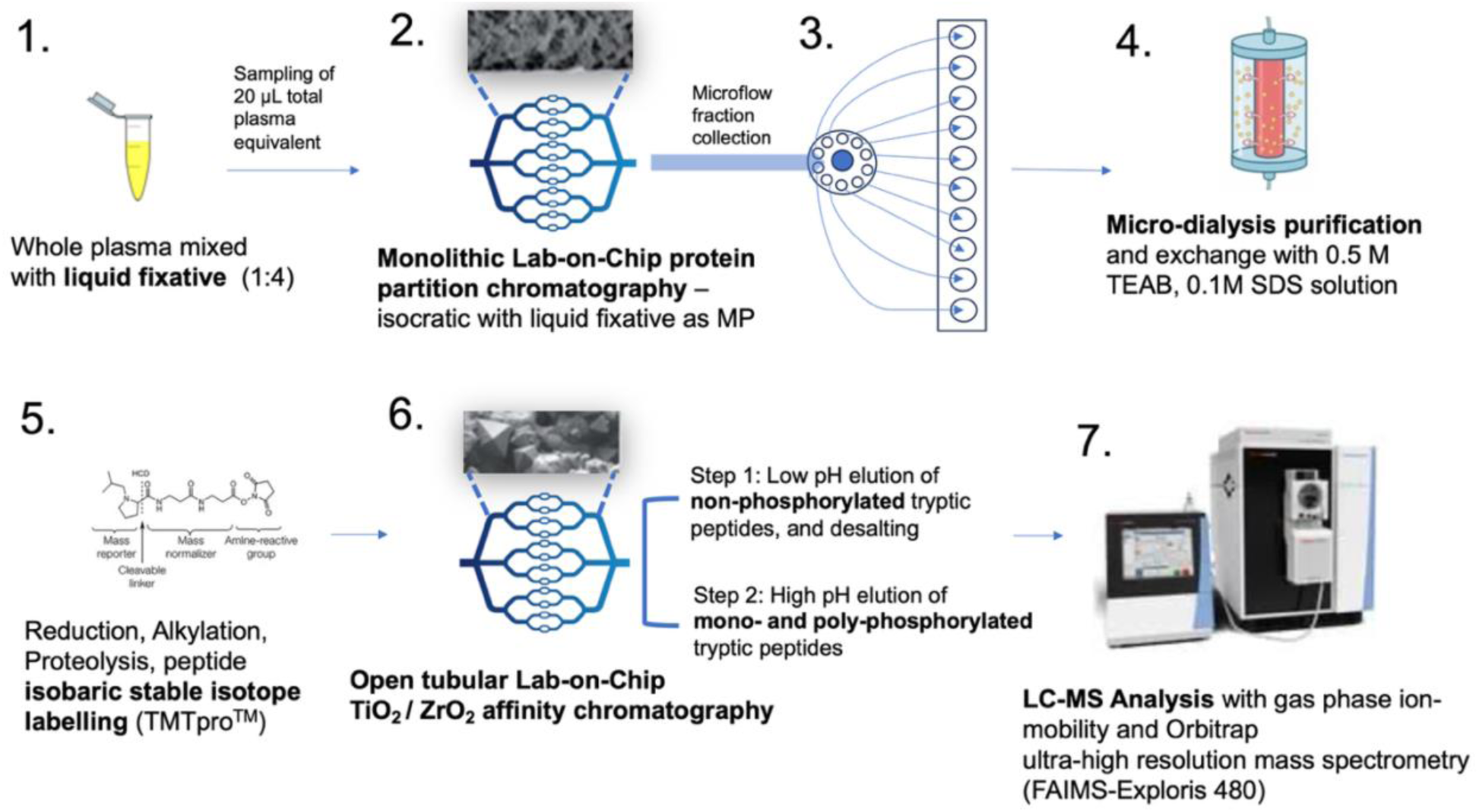
The basic steps of the Biofluid Total Analytic System (BioTAS) proteomics platform applied to study.

### Proteomics Data Processing

Unprocessed raw files were submitted to Proteome Discoverer 2.5 for target decoy search using Sequest. The UniProtKB homo sapiens database which comprised 20,159 entries (release date January 2025) was utilized. The search allowed for up to two missed cleavages, a precursor mass tolerance of 10ppm, a minimum peptide length of six and a maximum of two variable (one equal) modifications of; oxidation (M), deamidation (N, Q), or phosphorylation (S, T, Y). Methylthio (C) and TMT (K, peptide N-terminus) were set as fixed modifications. Peptide and protein FDRs were set at 0.01 (strict) and 0.05 (relaxed), percent co- isolation excluding peptides from quantitation was set at 20, with peptide confidence at least medium, lower confidence peptides excluded. Reporter ion abundances from unique peptides only were taken into consideration for the quantitation of the respective protein. Such peptide evaluation criteria have previously been reported by the authors (20, 23, 24, 26–29, 31, 32) and complied with the Proteome Discoverer PSM with peptide validation pipeline.

### Biostatistics

To identify differentially abundant proteins (DAPs) in patients treated with semaglutide, the respective protein abundance ratio for each analyzed protein was considered. Ratios were log_2_-transformed to normalize their distribution. A one-sample T-Test (paired) using the two-stage Benjamini Yekutieli Krieger step up method for multiple hypothesis testing FDR correction of the p-value was performed (33). The threshold of significance was set at *q<0.05*. Statistical power was determined with the pwr.t.test() function statistical package in R (34). The same procedure was applied to identify the differentially abundant phosphoproteins (DApPs).

### Key outcome

This study achieved an extremely high reported depth-and-breath in quantitative proteome and phosphoproteome coverage from an equivalent of only 20µL whole plasma equivalent from each patient. Namely, a total of 13,173 proteins and 25,578 phosphopeptides (Supplementary Table 4-6) were fully profiled (*q≦ 0.05*). Of these, 1,040 were measured as differentially abundant proteins (DAPs) and 1,064 were measured as differentially abundant phosphoproteins (DApPs) with an absolute log_2_-fold-change of ≧ 0.5, CV ≦ 15%, and q*≦ 0.05* (Supplementary Tables 4 - 7). Accounting for the multiplex intra-person pre- and post- treatment effects clinical design, these metrics yielded a statistical power ≧ 0.7 with the pwr.t.test() function (34) at standard deviation 0.77 and a fold change of 2 at the alpha significance level of 0.05, despite the small sample size. Protein abundance ratios were log_2_ transformed and shown as volcano plots for the total proteome abundance (Figure 2A) and phosphoproteome (Figure 2B). The x-axis indicates log_2_ fold change values while the y-axis indicates the adjusted negative log_10_ q-values (FDR step up procedure, 33). Proteins below a 0.05 q-value cutoff are shown in black while upregulated and downregulated proteins are shown in red and blue, respectively for the DAPs and sky blue and orange, respectively for the DApPs. Proteins with an absolute log_2_ fold change value > 1.5 are labeled (Figure 2A). Protein abundance values were log_2_ transformed and subjected to principal component analysis. The variance explained by each principal component is shown in the axes. Coloration indicates groups for samples with purple for semaglutide, green for baseline, and black for placebo for the proteome (Figure 2C), and the phosphoproteome (Figure 2D). in both cases, PCA could distinguish between semaglutide post treatment and baseline individuals as well as the placebo samples, albeit to a lesser extent for the phosphoproteome. Protein abundance values were log_2_ transformed and subset to the top 100 proteins by absolute log_2_ ratio between baseline and Semaglutide. A heatmap was generated using z-transformed log_2_ transformed abundance values with blue and red indicating a negative and positive z-scores respectively (Figure 2E). Phosphoprotein abundance values were log_2_ transformed and subset to the top 100 phosphoproteins by absolute log_2_ ratio between baseline and Semaglutide. A heatmap was generated using z-transformed log_2_ transformed abundance values with skyblue and orange indicating a negative and positive z-scores respectively (Figure 2F). Protein/phosphoprotein and sample dendrograms were generated using hierarchical clustering with a Euclidean distance metric. The top line indicates which samples are baseline (Green), treated with Semaglutide (purple), and placebo (grey) (Figure 2E and 2F). Additional images of PC3 vs. PC2 and PC3 vs. PC1 are included along with their total percent of variation are shown in the Supplementary Figures. The mass spectrometry proteomics data have been deposited to the ProteomeXchange Consortium via the PRIDE partner repository with the dataset identifier PXD063933.

**Figure 2.**
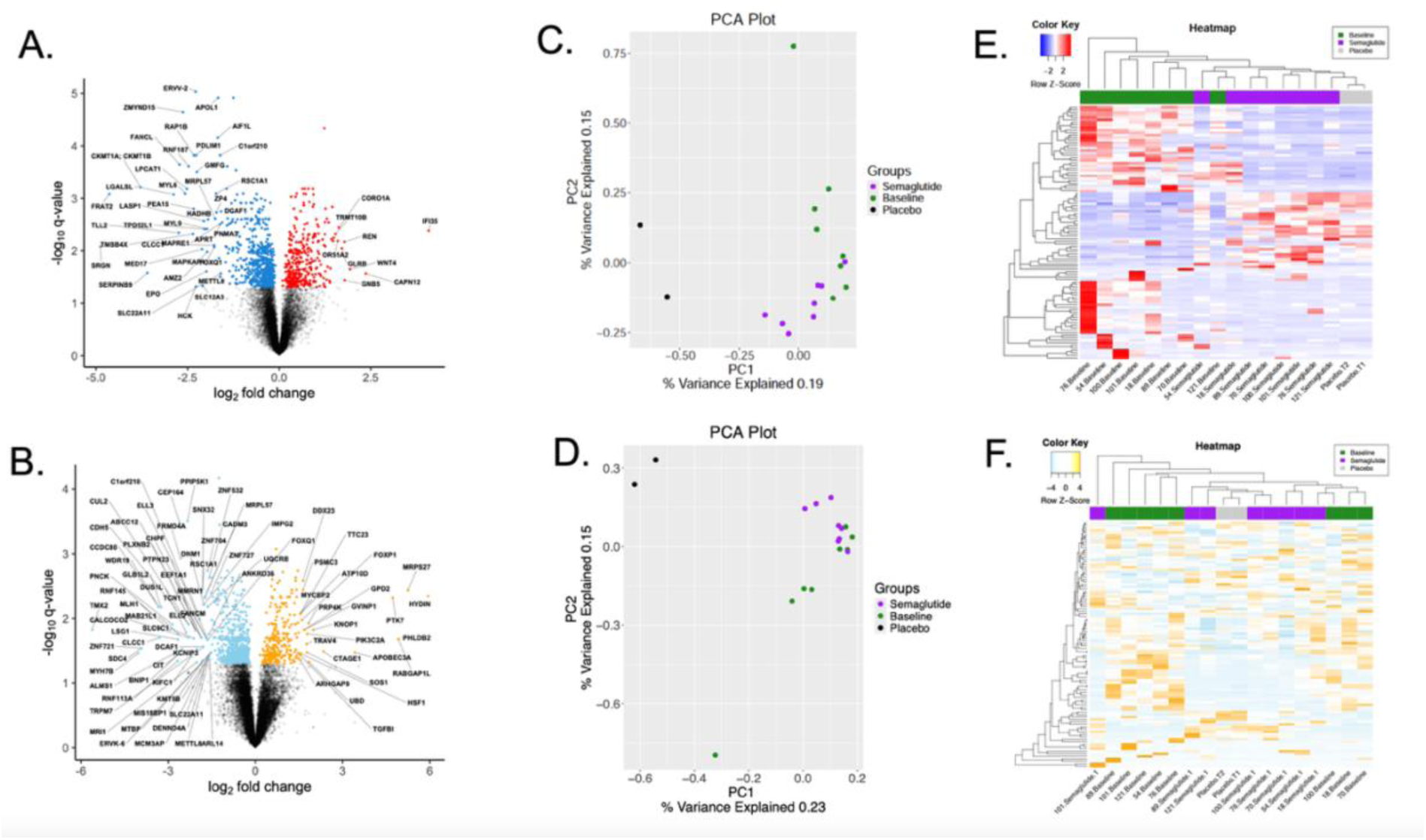
Protein abundance (A) and phosphoproteins (B) ratios were log2 transformed and shown in a volcano plot. The x-axis indicates log2 fold change values while the y-axis indicates the adjusted negative log10 q-values (FDR step up procedure, 33). Proteins below a 0.05 q-value cutoff are shown in black while upregulated and downregulated proteins are shown in red and blue respectively. Proteins with an absolute log2 fold change value > 1.5 are labeled. Protein abundance values were log2 transformed and subjected to principal component analysis (C. and D.). The Variance explained by each principal component is shown in the axes. Coloration indicates groups for samples with purple for semaglutide, green for Baseline, and black for placebo. Protein abundance (E) values were log2 transformed and subset to the top 100 proteins by absolute log2 ratio between baseline and semaglutide. A heatmap was generated using z-transformed log2 transformed abundance values with blue and red indicating a negative and positive z-scores respectively. Protein and sample dendrograms were generating using hierarchical clustering with a Euclidean distance metric. The top line indicates which samples are baseline (Green), treated with semaglutide (purple), and placebo (grey).

To examine the linear dynamic range of the plasma proteome and phosphoproteome profiles for this study we used the well curated PaxDB as the database of choice due to their methodology for estimating protein abundance data (46–48). PaxDB is a comprehensive compendium of plasma or serum proteomics and phosphoproteomics datasets generated with diverse methods and protocols and multiple types of mass spectrometry instrumental platforms such as those based on quadrupole-time of flight, quadrupole-linear or quadratic ions trap, and hybrid and tribrid Orbitrap designs (45). These diverse datasets are mapped onto a common namespace, and processed using their standardized spectral counting pipeline. These datasets also include those generated with more specialized plasma or serum exosome isolation and enrichment protocols followed by their mass spectrometry-based analysis for their proteome content (49). Such a compilation for these datasets allows for a more robust estimate of protein abundance data free from individual study bias. Protein abundance datasets in PaxDb are re-scaled to a common abundance metric (’parts per million’), and also ranked via a universally applicable, albeit somewhat indirect quality score. For the re-scaling, the datasets are first parsed or processed such that the data reflect proportional abundances of whole proteins (i.e. proportionality to counts of complete, individual proteins, not to molecular weights, protein volumes, or digested peptides). Overall, the PaxDB resulted in a robust and well curated total compilation of about 6,500 human plasma and serum proteins observed to date. The application of PaxDB to the total proteome and phosphoproteome for this study generated a linear dynamic range curve spanning over 11-orders of magnitude (Figure 3A). Example protein names were indicated. Accounting for their molecular weights, spectral counts, and part-per-million abundance levels, the native protein concentration range encompassed by the observed dynamic range curve span from mg/L down to low fg/L. Of additional interest, > 85% of the PaxDB database was profiled in this study. The remaining approx. 9,000 proteins and phosphoproteins for this study constituted entirely novel plasma matrix observations.

### Exosomal protein annotation

To identify which of the PaxDB proteins also profiled in this study were of exosomal origin, the ExoCarta database was employed (50,51). ExoCarta is a web-based database and compendium of proteins (along with lipids, RNA and DNA) experimentally found in exosomes and other lipid microvessicles from published and unpublished studies. In addition to providing detailed information of exosomal cargo composition, ExoCarta also provides valuable information for their biological role, tissue or cell type of origin, along with their experimental methods used to identify them (50,51). Cross examination with the ExoCarta database showed that ∼70% of the proteins and phosphoproteins of this study, which were congruent with the PaxDB database, were of exosomal origin. Figure 3A included annotation of proteins and phosphoproteins profiled or differentially abundant that were also exosome derived. Accounting for the total DAPs and DApP profiled from this study, Figure 3B provides a Venn diagram description of their origin as novel protein or phosphoproteins observed directly or derived from exosomes in plasma. These are deemed significant findings given the low abundance of exosome derived proteins in plasma along with their more specialized and technically challenging methods required to isolate them from larger serum or plasma starting volumes typically ranging from 500 μL – 10 mL (49). By contrast, the proteomics platform used for this study, achieved an unprecedented level of plasma exosome proteome/phosphoproteome coverage from only 20 μL plasma equivalent volume starting material per clinical specimen. Achieving such a level in exosome proteomics / phosphoproteomics coverage was requisite to the discovery of clinically and biological relevant protein signatures, given the well-established role of plasma exosomes for the communication and regulation of disease affected tissue compartments and their response to treatment (12–16).

**Figure 3.**
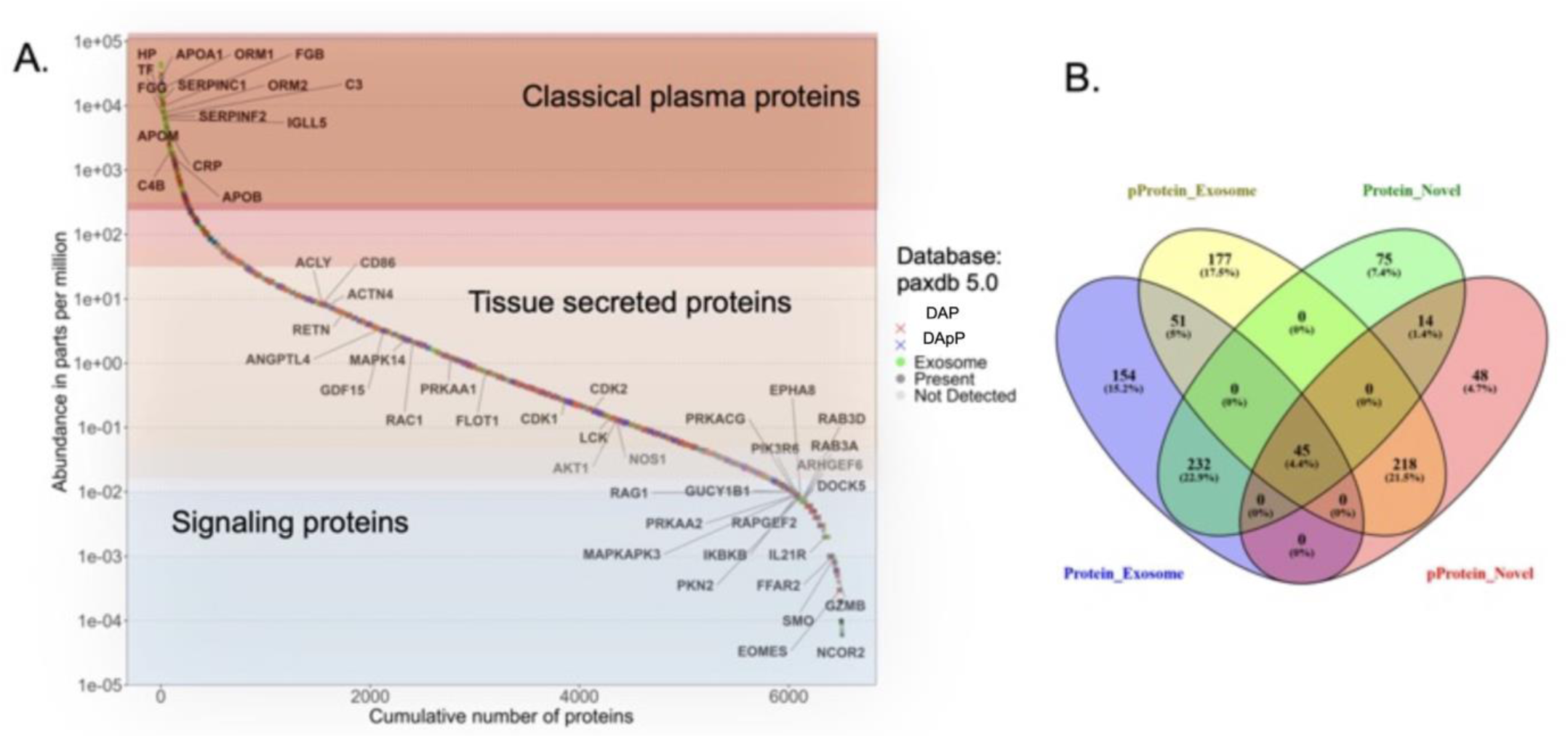
Differentially abundant proteins were mapped onto the PaxDB database (A). The y-axis shows protein abundance in parts per million (ppm), and the x-axis shows proteins ranked by ppm. Three categories are highlighted: signaling proteins, secreted proteins, and classical plasma proteins. Differential proteins are marked with a red “X,” phosphorylated proteins with a blue “X,” overlapping proteins between our dataset and PaxDB in dark gray, and proteins present only in PaxDB in light gray. Differentially abundant and phosphorylated proteins were further classified as significant exosome proteins/phosphoproteins or significant novel proteins/phosphoproteins, with “novel” indicating absence from the PaxDB plasma dataset and shown in a Venn diagram (B).

### PROMINIA Computational Biology

The PROMINIA computational biology suite was developed to provide an intelligent inference and causal pathway deconvolution of the experimentally observed proteins and phosphoproteins. In particular, PROMINIA integrates the DAPs and DApPs (with specific phosphosites) with the respective transcription factors and kinases that regulate them. Its basic operating tenets are described below.

### Quantitative proteomics

**(DAPs)** provides a descriptive snapshot of the abundant proteins in terms of their cellular topology, molecular and biological functionality. These properties are accounted for in their pathway enrichment.

### Quantitative phosphoproteomics

(DApPs) provides a dynamic snapshot for active signaling pathways affected by a disease state and its response to treatment. Quantification of the phosphorylation state of a given protein signifies its functional activity, primarily driven by upstream kinases implicated in phosphorylating specific protein substrates.

### Determination of kinase activity

PROMINIA facilitates the use of quantitative phosphoproteomics evidence (DApPs), to infer the activity of upstream or downstream kinases implicated with the phosphorylation of specific target proteins that also accounts for specific S, T or Y amino acid residues measured. This approach provides a robust inference of specific kinase activity guided by the quantitative phosphoproteomics-based evidence. Such a functional association strengthens the causal link to a specific signaling pathway or protein-to-protein interaction network.

### Contextualizing quantitative proteomics (DAPs) with transcription factors (TFs)

TFs control gene expression, which in turn affect protein abundance. By inferring TF to the observed DAPs, causal links between gene regulation and protein expression could be achieved. TFs have also been implicated in the intracellular regulation for the secretion of both soluble and exosomal proteins in the systemic circulation.

### Z-score quantitation for pathway direction of change (induction or inhibition)

To determine overall direction of change for each pathway the following equation:

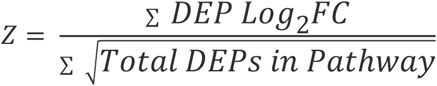

For this calculation, the numerator was the sum of all statistically significant fold change and the denominator is the square root of the total number of DAPs found in the pathway. This gives an overall measure of up- (positive z-score) or down-regulation (negative z-score) of the pathway.

These components are then mapped to specific canonical pathways either induced or inhibited, as determined by the overall z-score, with high statistical confidence (Chi-squared test, FDR corrected, *p<0.001*). Integrating these distinct but mutually reinforcing layers of biological information creates a more holistic and dynamic picture of disease pathway signaling. Consequently, such a strategy can support a deeper understanding of cause-and-effect relationships rather than correlations that typify mainstream biomarker discovery studies. Figure 4 provides an overview of the PROMINIA computational biology pipeline used to interrogate the DAPs and DApPs to ultimately provide a deeper molecular understanding of semaglutide treatment to patients with atherosclerosis and T2D.

**Figure 4.**
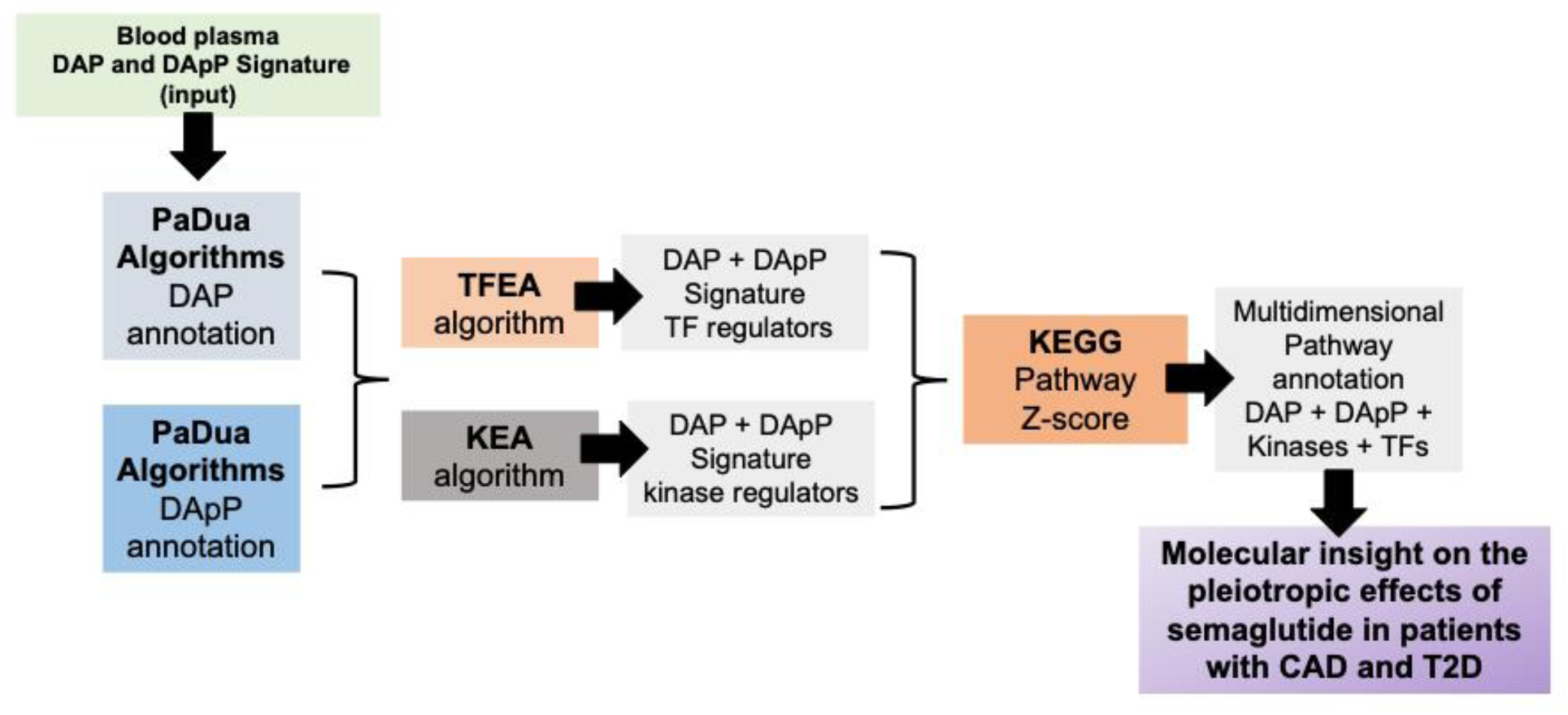
The modular components that constitute the PROMINIA computational biology pipeline.

More detailed descriptions of the PROMINIA components are provided below:

### Phosphoproteomics data processing

The interpretation of the differentially abundant phosphoproteins (DApP) was facilitated with the PaDuA Python software package specifically geared for high-throughput, mass spectrometry-based data sets (37). PaDuA contextualizes and correlates the phosphoproteomics data as cellular signaling networks making use of library processes and filters from large-scale protein phosphorylation studies utilizing mass spectrometry-based approaches.

### Transcription Factor Enrichment Analysis of DAPs

DAPs (FDR corrected p-value ≤ 0.05 and absolute log_2_FC values ≥ 0.5) were inserted into the ChEA3 algorithm available at (https://maayanlab.cloud/chea3/). CHEA3 performs transcription factor enrichment analysis using ChIP-Seq experiments from ENCODE, ReMap, and individual well reviewed publications (38). Co-expression of TFs with other proteins is done by examining them against thousands of proteins lists submitted to the tool Enrichr. A fisher’s exact test is performed (background of 20k proteins and/or genes) to compare and closely associate the input protein set to the TFs that target them. The MeanRank algorithms that account for the mean value of the rankings from datasets used and a low value suggests a broad agreement among multiple databases regarding the importance of this item given the gene/protein list. In the mean rank method, the implicated TFs are shown in tabular form in each of the affected signaling pathways (Table 1).

**Table 1.** Biologically relevant pathways. KEGG pathways were selected based on their biological relevance to cardiac function, liver fibrosis, insulin signaling, inflammation, and diabetes-associated comorbidities. The table reports count of differentially abundant proteins, differentially abundant phosphorylated proteins, predicted transcription factors, and predicted kinases. Z-scores were derived by summing the log fold changes and dividing by the square root of the number of differentially abundant proteins/phosphorylated proteins. P-values were determined using chi-squared testing with Benjamini–Hochberg (B&H) FDR correction. Pathway specific listings for the DAPs, DApPs, TFs, and kinases along with the annotated Kegg-pathway for each are listed in the Supplementary Tables 8-12.

| Kegg Pathway | Diff. Abund.<br>Proteins (DAP) | Diff. Exp.<br>Phosphoproteins<br>(DAPP) | Transcription<br>Factors | Kinases | z-score<br>DAPs | z-score<br>DAPPs | FDR-corrected q-value |
| --- | --- | --- | --- | --- | --- | --- | --- |
| AGE-RAGE signaling | 10 | 10 | 6 | 21 | -0.34 | -1.82 | $6.54 \times 10^{-6}$ |
| Insulin signaling | 15 | 7 | 1 | 27 | 0.48 | -0.88 | $4.80 \times 10^{-4}$ |
| Lipid and<br>atherosclerosis | 35 | 9 | 12 | 23 | -1.41 | -1.64 | $2.09 \times 10^{-8}$ |
| Non-alcoholic fatty<br>liver disease | 26 | 6 | 8 | 15 | -1.37 | -1.61 | $2.16 \times 10^{-8}$ |
| Diabetic<br>cardiomyopathy | 22 | 9 | 2 | 21 | -1.32 | -2.15 | $7.27 \times 10^{-3}$ |

### Kinase Enrichment Analysis (KEA3)

DAPs (FDR corrected p-value ≤ 0.05 and absolute log_2_FC values ≥ 0.5) were inserted into the KEA3 algorithm available from (https://maayanlab.cloud/kea3) to determine if this protein set is enriched with proteins known to interact with kinases. KEA3 uses a gene-set library with kinases and their interacting sets of proteins curated from well published studies against *in vivo* models and those involving human tissue or primary cells. The KEA algorithm also accounts for specific phosphosites experimentally observed. The algorithm “Mean rank” is the mean value of the rankings from datasets used by KEA and a low value suggests a broad agreement among multiple databases regarding the importance of this kinase given the protein list (39,40). The implicated kinases are shown in tabular form in each of the affected signaling pathways (Table 1). These predicted kinases were assessed for relationship to conditions of interest. These results indicate a heavy presence of kinases related to these conditions and could be of utility as treatment targets.

### Pathway annotation and integration

Kegg pathways are annotated with the observed differentially abundant proteins (DAPs) and phosphoproteins (DApPs) along with the predicted transcription factors (TFs), and kinases (Ks) affected to better illustrate their synergistic interaction based on an up-to-date and curated database of published studies using *in vivo* models and human derived tissue or primary cells. Images were generated using the KEGG Pathway analysis tool (https://www.genome.jp/kegg/pathway.html). These pathways were subcategorized based on primary pathophysiology affected. It must be noted however, that such pathways are functionally interdependent and mutually reinforcing given their association with fundamental biochemical and molecular biology processes known to drive cardiometabolic spectrum comorbidities.

## RESULTS

Table 1 is a compilation for the number of DAPs, DApPs, enriched TFs and kinases associated with each Kegg canonical pathway. Their itemized listings are provided in the Supplementary Tables 8-12. As a means to illustrate the biomedical utility of the BioTAS platform, an overview of the canonical pathway evidence that also support the deeper molecular understanding of the reported cardiac CT imaging biomarker and blood biochemistry profiling observations to cardiovascular health (6,7), and liver fat (8) from the same STOP trial is presented.

### Inhibition of AGE-RAGE signaling pathway

In plasma specimens obtained from participants enrolled in the Semaglutide Treatment Effect in People with Obesity (STOP) trial (7), semaglutide therapy was associated with coordinated suppression of the advanced glycation end product–receptor for AGE (AGE–RAGE) signaling pathway compared with placebo (**Figure 5, Supplementary Table 8**). Differential abundance analysis demonstrated pathway-level inhibition at both the total protein (DAP z-score = −0.34) and phosphoprotein levels (DApP z-score = −1.82), with a more pronounced effect observed for phosphorylation-dependent signaling events. In the STOP trial, semaglutide significantly improved conventional glycemic indices, including fasting plasma glucose and hemoglobin A1c, relative to placebo.

**Figure 5.**
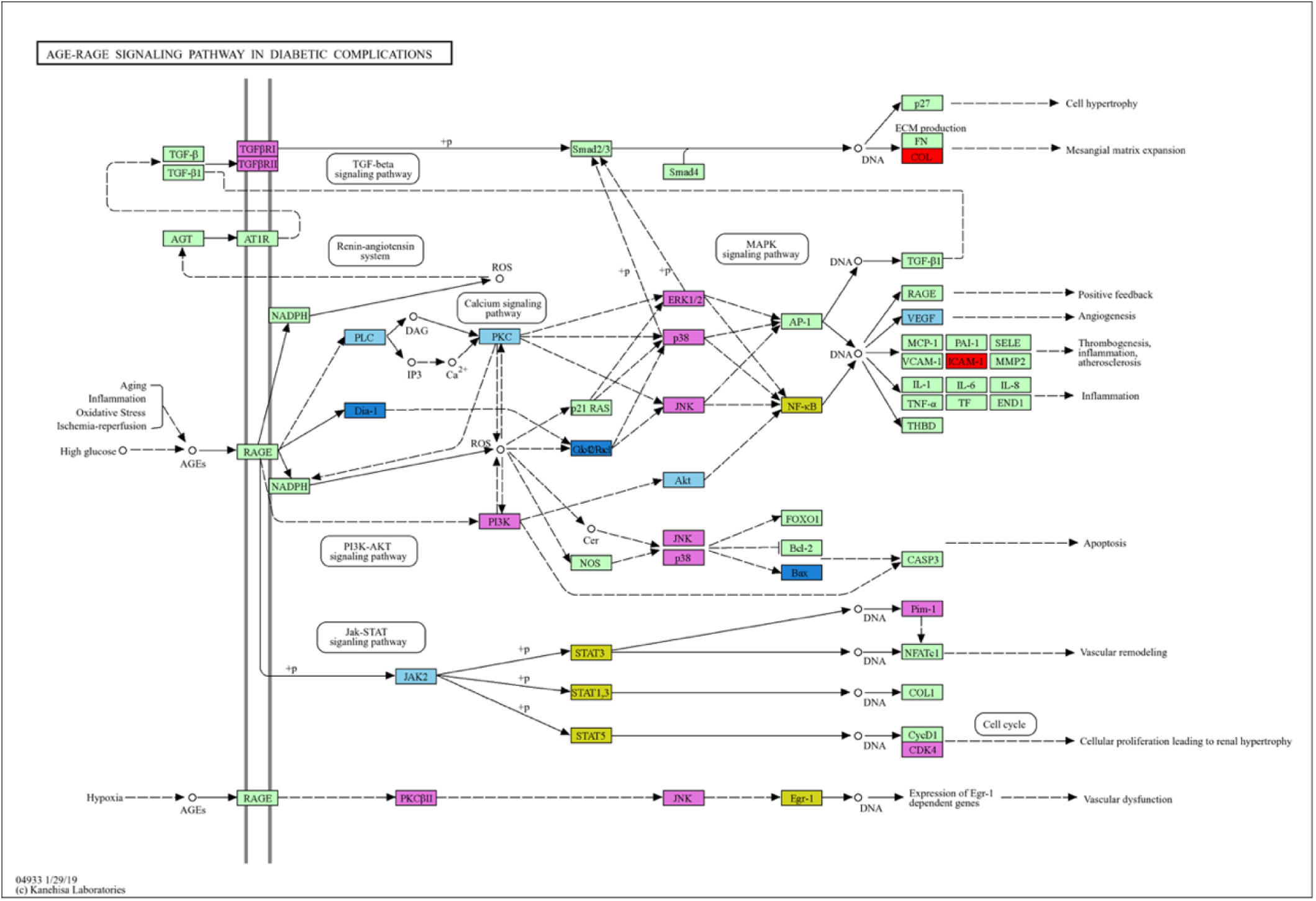
AGE-RAGE Signaling pathway inhibited by Semaglutide (DEP, z-score=-0.34; DApP, z-score=-1.82). Pathway mapping was performed by overlaying differentially expressed proteins, predicted transcription factors, and predicted kinases onto KEGG pathways and visualized by category: upregulated proteins (red), downregulated proteins (blue), upregulated phosphoproteins (orange), downregulated phosphoproteins (sky blue), predicted transcription factors (yellow), and predicted kinases (purple). Statistical significance was determined using chi-squared testing with Benjamini–Hochberg false discovery rate (FDR) correction (q = 6.54 × 10^-6^). Predicted transcription factors and kinases were identified through overrepresentation enrichment analysis of the differentially expressed proteins.

AGE–RAGE signaling is a central mediator linking chronic hyperglycemia to vascular inflammation, oxidative stress, endothelial dysfunction, and accelerated atherogenesis in type 2 diabetes (52–54,60). RAGE functions as a multiligand pattern-recognition receptor that amplifies immune and inflammatory responses (52), and AGE accumulation directly contributes to diabetic vascular injury and coronary artery disease progression (53). At the cellular level, AGE engagement of RAGE activates redox-sensitive signaling cascades and NF-κB–dependent transcriptional programs that sustain oxidative stress and inflammatory gene expression (54,60).

Consistent with this biology, semaglutide-treated participants demonstrated suppression of both proximal and distal components of the AGE–RAGE axis. At the proximal interface, diaphanous-related formin 1 (DIAPH1)—an obligate cytoplasmic binding partner required for RAGE signal transduction—was significantly reduced (55). Interaction between the RAGE cytoplasmic domain and DIAPH1 is required for Rac1- and Cdc42-mediated cytoskeletal activation and downstream inflammatory signaling (55). Pharmacologic inhibition of ligand-stimulated RAGE–DIAPH1 signal transduction attenuates inflammatory and vascular responses (56), and recent translational work further implicates the RAGE/DIAPH1 axis as a critical driver of atherosclerosis progression (57).

Importantly, the more pronounced suppression at the phosphoprotein level reflected coordinated inhibition of associated kinase networks downstream of RAGE engagement. Reduced phosphorylation of protein kinase C (PKC), phospholipase C (PLC), and Janus kinase 2 (JAK2) was observed, consistent with attenuation of kinase-driven inflammatory flux (64–66). PKC activation is a well-established contributor to diabetic vascular complications (64), and AGE–RAGE signaling activates JAK2/STAT pathways in vascular and inflammatory cells (65,66). Experimental inhibition of JAK2 phosphorylation attenuates pathological cardiac hypertrophy and pressure-overload–induced remodeling (58,59), underscoring the cardiovascular relevance of this signaling node. The reduction in JAK2 phosphorylation observed in semaglutide-treated participants therefore suggests broader suppression of pro-hypertrophic and pro-inflammatory kinase cascades implicated in diabetic cardiomyopathy and coronary artery disease.

Concomitant inhibition of vascular endothelial growth factor (VEGF) pathway phosphorylation was also observed. VEGF signaling plays a central role in vascular permeability, angiogenesis, and plaque neovascularization (61). AGE–RAGE–dependent PKC and NF-κB activation induces VEGF expression (53,54), linking hyperglycemia to microvascular dysfunction and plaque instability. Suppression of VEGF pathway phosphorylation in this cohort indicates attenuation of downstream angiogenic and permeability signaling, further supporting vascular stabilization.

Database interrogation revealed that a substantial proportion of differentially abundant AGE–RAGE–associated proteins and phosphoproteins were of exosomal origin, suggesting modulation of extracellular vesicle–mediated inflammatory propagation (12–16). This systems-level inhibition of extracellular inflammatory signaling provides additional mechanistic context for the cardiovascular benefit observed with GLP-1 receptor agonists in large-scale outcome trials. Notably, semaglutide significantly reduced major adverse cardiovascular events in SUSTAIN-6 (62), and dulaglutide demonstrated cardiovascular benefit in REWIND (63).

Importantly, while fasting glucose and HbA1c quantify systemic glycemic exposure, they do not directly capture the underlying inflammatory and kinase-driven signaling consequences of chronic hyperglycemia. In contrast, the integrated panel of AGE–RAGE–associated proteins and phosphorylation-dependent kinase substrates reflects dynamic pathway activity and intracellular signaling flux. These phospho-resolved (including exact protein-wide phosphosite topology) protein markers therefore provide complementary and mechanistically informative indices beyond traditional metabolic measures. Such markers may serve as pharmacodynamic indicators of GLP-1 receptor agonist target engagement, enabling dose optimization and assessment of treatment frequency. Furthermore, they provide a translational framework for evaluating additive or synergistic effects when GLP-1 receptor agonists are combined with complementary small-molecule inhibitors targeting RAGE–DIAPH1 interaction, PKC isoforms, or JAK2/STAT signaling (55–59,64–66), as well as with lifestyle interventions known to reduce AGE burden (67,68).

#### Inhibition of Insulin Signaling Pathway

In plasma specimens obtained from participants enrolled in the STOP trial, semaglutide therapy was associated with coordinated remodeling of the insulin signaling pathway, as defined by integrated proteomic and phosphoproteomic profiling (**Figure 6, Supplementary Table 9**). Pathway enrichment analysis demonstrated a bidirectional regulatory pattern characterized by induction at the total protein level (DAP z-score = 0.48) and relative suppression at the phosphorylation level (DApP z-score = −0.88).

**Figure 6.**
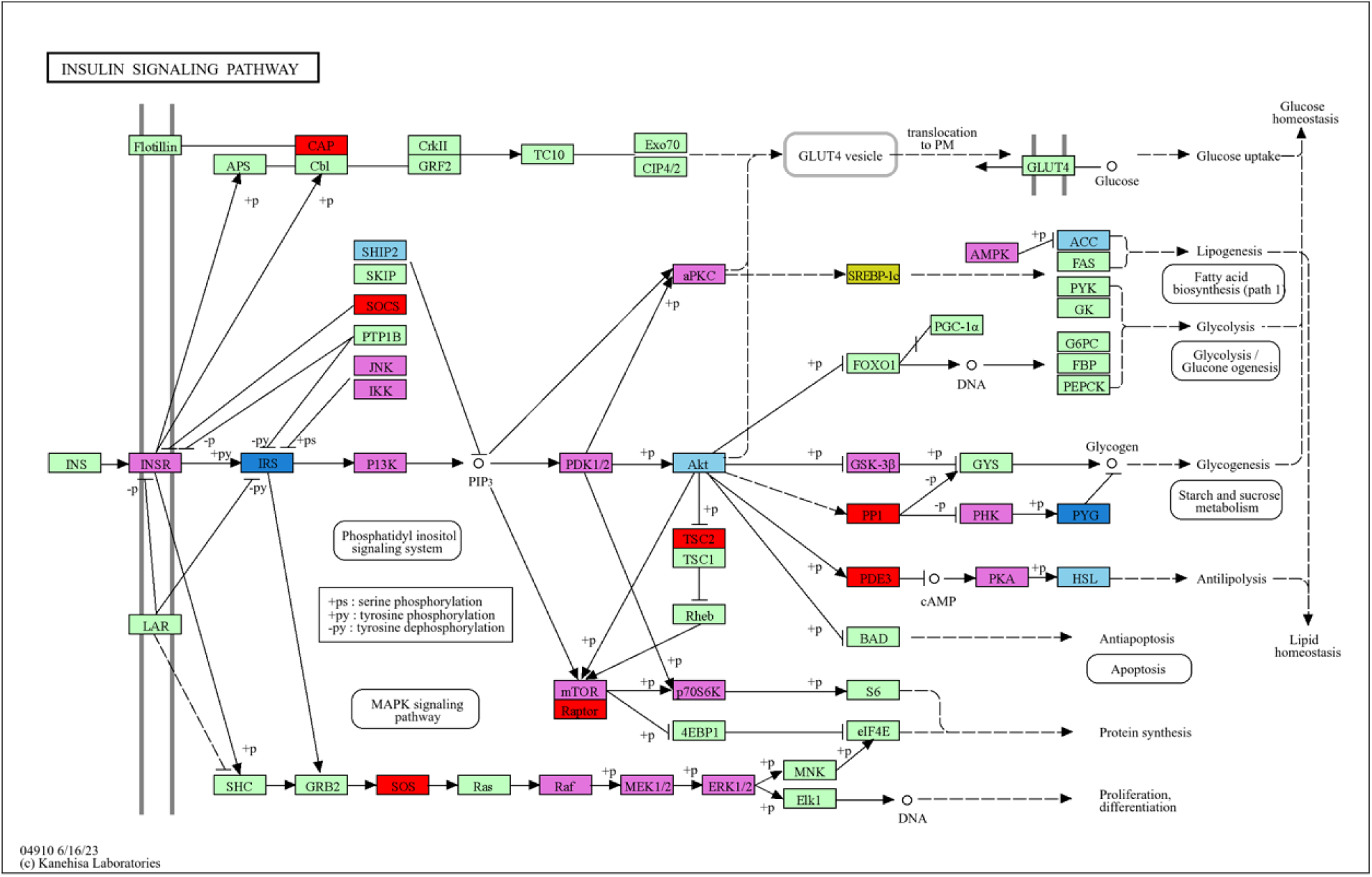
Insulin Signaling Pathway mapping was performed by overlaying differentially expressed proteins, predicted transcription factors, and predicted kinases onto KEGG pathways and visualized by category: upregulated proteins (red), downregulated proteins (blue), downregulated phosphoproteins (sky blue), predicted transcription factors (yellow), and predicted kinases (purple). Statistical significance was determined using chi-squared testing with Benjamini–Hochberg false discovery rate (FDR) correction (q = 4.8 × 10^-4^). Predicted transcription factors and kinases were identified through overrepresentation enrichment analysis of the differentially expressed proteins.

These molecular alterations occurred in parallel with significant reductions in fasting plasma glucose and hemoglobin A1c compared with placebo, confirming improved glycemic control in the semaglutide-treated group (6–8).

GLP-1 receptor agonists enhance pancreatic β-cell responsiveness and improve peripheral insulin sensitivity through modulation of PI3K–AKT signaling (69,70). As shown in Fig. 5, and consistent with this mechanism, semaglutide-treated participants demonstrated increased abundance of insulin receptor substrate-1 (IRS1) with concomitant reduction of IRS2. IRS1 upregulation is associated with enhanced phosphoinositide 3-kinase (PI3K) activation and increased generation of phosphatidylinositol-3,4,5-trisphosphate (PIP3), promoting downstream AKT signaling and glucose uptake (71). This effect was further supported by reduced phosphorylation of SH2-containing inositol 5-phosphatase 2 (SHIP2), a negative regulator of PI3K signaling (72–74). Experimental inhibition or deletion of SHIP2 enhances insulin sensitivity and improves glucose tolerance (72,73), consistent with the insulin-sensitizing signature observed in this cohort.

Downstream metabolic nodes were also favorably modulated. Increased abundance of protein phosphatase-1 (PP1) was observed, a key regulator that suppresses hepatic and skeletal muscle glycogenolysis. PP1 inhibits phosphorylase kinase (PhK), thereby reducing phosphorylation and activation of glycogen phosphorylase (PYG), ultimately limiting hepatic glucose output (75,76). This mechanism is particularly relevant under fasting or GLP-1RA–mediated fasting-mimetic states (77), aligning with the observed reduction in circulating glucose levels. Reduced phosphorylation of hormone-sensitive lipase (HSL) was also detected, consistent with suppression of excessive lipolysis and reduced free fatty acid flux—mechanisms known to improve insulin sensitivity in experimental models (78). In addition, inhibition of phosphorylated acetyl-CoA carboxylase (p-ACC) was observed, consistent with enhanced fatty acid oxidation and improved metabolic flexibility, paralleling metabolic effects reported with AMPK activation (79).

Integration with the ExoCarta database revealed that the majority of insulin signaling–related differentially abundant proteins and phosphoproteins were of exosomal origin, implicating endothelial cells, pancreatic β-cells, hepatic cells, adipocytes, skeletal muscle cells, and macrophages as contributors. This systems-level extracellular vesicle signature parallels the AGE–RAGE pathway findings (12–16,52–57,60,64–66) and supports a unified model in which semaglutide modulates intercellular inflammatory–metabolic communication networks central to cardiometabolic disease progression.

Importantly, remodeling of insulin signaling occurred concurrently with coordinated suppression of the AGE–RAGE axis in the same participants (52–57,60,64–66). AGE–RAGE activation impairs insulin receptor signaling through oxidative stress–mediated serine phosphorylation of IRS proteins and activation of stress kinases (53,54,60). The simultaneous attenuation of AGE–RAGE–dependent inflammatory signaling and restoration of insulin pathway phospho-dynamics therefore supports an integrated mechanistic foundation linking reduced oxidative stress to improved insulin sensitivity.

These molecular and phosphorylation-dependent changes were accompanied by favorable remodeling of ectopic fat depots. Semaglutide treatment significantly reduced epicardial adipose tissue volume in the STOP imaging sub study (6), a metabolically active visceral fat depot implicated in coronary inflammation. Serial cardiac CT imaging also demonstrated significant reductions in hepatic fat content compared with placebo (8), consistent with improved hepatic insulin sensitivity and reduced lipotoxic stress. Baseline coronary characterization of the STOP cohort by coronary computed tomography angiography further contextualized these findings within a population with established atherosclerotic risk (7).

Collectively, semaglutide-mediated reductions in blood glucose and HbA1c were accompanied by coordinated enhancement of insulin-sensitizing protein networks, suppression of maladaptive kinase phosphorylation events, reduction in epicardial and hepatic fat depots, and parallel inhibition of AGE–RAGE–dependent inflammatory signaling. This integrated molecular and imaging signature provides mechanistic evidence linking GLP-1 receptor agonist therapy to improved insulin sensitivity, reduced ectopic fat burden, attenuation of inflammatory–metabolic cross-talk, and modification of cardiometabolic risk substrates in type 2 diabetes.

#### Lipid and Atherosclerosis pathway

Consistent with prior cardiovascular outcomes trials demonstrating macrovascular benefit with GLP-1 receptor agonists (62,63), semaglutide treatment in the STOP cohort was associated with coordinated inhibition of lipid-driven atherogenic signaling. Cardiac computed tomography–derived imaging biomarkers demonstrated significant reductions in epicardial adipose tissue (EAT) volume and favorable changes in EAT density in semaglutide-treated participants compared with placebo (6). Given that EAT is a metabolically active visceral fat depot contiguous with the coronary arteries, pathological expansion is linked to hypoxia, increased lipolytic flux, and secretion of pro-inflammatory adipokines and cytokines, thereby promoting local oxidative stress and coronary atherogenesis.

Complementing these imaging data, application of the Proteas BioTAS platform to plasma samples from the same STOP cohort (6–8) identified coordinated inhibition of the lipid and atherosclerosis pathway (**Figure 7; Supplementary Table 10**), with significant reductions at both the total protein (DAP z-score = −1.41) and phosphoprotein (DApP z-score = −1.64) levels. This pathway reflected convergent modulation of multiple interconnected signaling axes—including AGE–RAGE (52–57,60), Toll-like receptor (TLR), PI3K–AKT (69–71), tumor necrosis factor-α (TNFα), and p53—integrating lipid-derived stimuli such as very-low-density lipoprotein (VLDL), oxidized and glycated LDL (oxLDL/AGE-LDL), triglycerides, and apolipoprotein B–containing particles.

**Figure 7.**
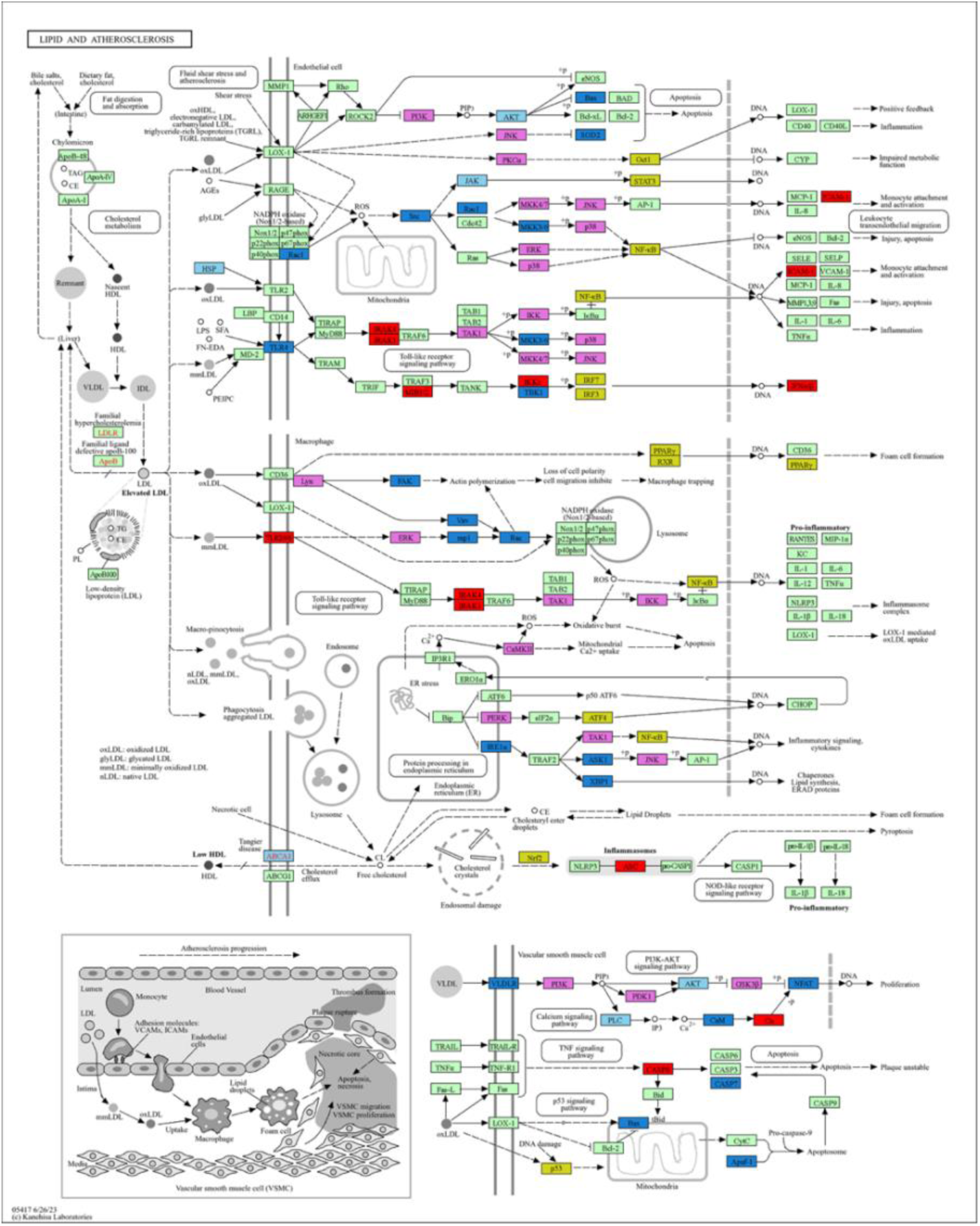
Lipid and atherosclerosis pathway mapping (DEP *z-score=-1.41;* DApP *z-score=-1.64*) was performed by overlaying differentially expressed proteins, predicted transcription factors, and predicted kinases onto KEGG pathways and visualized by category: upregulated proteins (red), downregulated proteins (blue), upregulated phosphoproteins (orange), downregulated phosphoproteins (sky blue), predicted transcription factors (yellow), and predicted kinases (purple). Statistical significance was determined using chi-squared testing with Benjamini–Hochberg false discovery rate (FDR) correction (q = 2.09 × 10^-8^). Predicted transcription factors and kinases were identified through overrepresentation enrichment analysis of the differentially expressed proteins.

TNFα-dependent inflammatory signaling is a central driver of plaque progression and destabilization (80,81). Experimental disruption of TNFα signaling attenuates atherosclerosis in ApoE-deficient models (83), and clinical anti-inflammatory targeting reduces cardiovascular events (82). Likewise, oxidized LDL promotes endothelial activation, foam cell formation, and apoptosis (85,86), processes amplified in diabetes by AGE-modified lipoproteins engaging RAGE (87). Macrophage-driven inflammatory amplification within the plaque further links modified lipoproteins to TNFα and NF-κB activation (84,88).

The p53 pathway also plays a critical role in vascular cell apoptosis, plaque instability, and DNA damage–associated atherogenesis (89,90). Lipid-induced oxidative stress and mitochondrial dysfunction activate p53-dependent pathways that contribute to necrotic core expansion and plaque vulnerability (89). The coordinated suppression of p53-associated phosphoproteins observed in semaglutide-treated participants is therefore consistent with attenuation of oxidative DNA damage signaling within the atherosclerotic milieu.

Triglyceride-rich lipoproteins and VLDL remnants are independently associated with cardiovascular risk (91), and apolipoprotein B–containing particles are causally linked to atherogenesis (92,93). VLDL-derived remnants and small dense LDL species promote endothelial dysfunction and inflammatory activation (94). The inhibition of lipid- and apolipoprotein-associated signaling nodes observed here aligns with attenuation of these pro-atherogenic lipid pathways.

Importantly, AGE–RAGE signaling functions as a mechanistic bridge linking dyslipidemia, oxidative stress, and chronic vascular inflammation in diabetes (52–54,60,87). Modified lipoproteins—including glycated LDL—engage RAGE to amplify NF-κB–dependent transcription and endothelial adhesion molecule expression, accelerating monocyte recruitment and plaque progression. The concurrent suppression of AGE–RAGE, insulin resistance–associated kinase pathways (69–74), and lipid-driven inflammatory cascades supports a unified model in which semaglutide disrupts pathogenic lipid–glucose cross-talk central to coronary artery disease progression.

These molecular findings complement the imaging evidence of reduced epicardial and hepatic fat depots (6,8) and align with macrovascular event reduction observed in SUSTAIN-6 and REWIND (62,63). Collectively, semaglutide therapy in the STOP trial was associated with coordinated suppression of lipid-driven inflammatory and apoptotic signaling at both protein and phosphorylation levels, reduction of pro-atherogenic visceral fat depots, and parallel inhibition of AGE–RAGE and insulin resistance pathways. This integrated systems-level signature substantiates a mechanistic framework in which GLP-1 receptor activation attenuates dyslipidemia-induced oxidative stress, TNFα-mediated inflammation, p53-associated vascular injury, and plaque progression in patients with type 2 diabetes.

### Coordinated Inhibition of the Non-Alcoholic Fatty Liver Disease (NAFLD) Pathway

As a prespecified secondary imaging outcome of the STOP trial, hepatic steatosis was assessed using serial non-contrast cardiac CT imaging (8). Semaglutide-treated participants demonstrated a significant reduction in liver fat compared with placebo, consistent with emerging evidence supporting GLP-1 receptor agonists as therapeutic agents for NAFLD and metabolic dysfunction–associated steatotic liver disease (95,96). Given the close pathophysiologic coupling between hepatic steatosis, insulin resistance, dyslipidemia, and systemic inflammation, these imaging findings extend the integrated cardiometabolic remodeling observed across AGE–RAGE (52–57,60), insulin signaling (69–74), and lipid–atherosclerosis pathways (80–94).

Integration with the ExoCarta database demonstrated that the majority of differentially abundant proteins and phosphoproteins mapped to the NAFLD pathway were of exosomal origin. This observation aligns with prior studies demonstrating increased hepatic exosome secretion in NAFLD, contributing to macrophage activation, systemic inflammation, and inter-organ metabolic dysfunction (99,100). These findings parallel the exosome-associated signatures observed in the AGE–RAGE and insulin pathway analyses, reinforcing a systems-level model of extracellular vesicle–mediated inflammatory–metabolic cross-talk.

At the mechanistic level, inhibition of TNFα-associated signaling is particularly relevant, as TNFα promotes hepatic insulin resistance, steatosis, hepatocellular injury, and progression to steatohepatitis (101,102). Concurrent enhancement of insulin pathway components (69–74) supports improved hepatic glucose handling and reduced de novo lipogenesis. Modulation of PPAR signaling further substantiates metabolic remodeling, given the regulatory role of PPARα and PPARγ in fatty acid oxidation, lipid storage, and adipocytokine balance (103,104).

Semaglutide-mediated inhibition of upstream RAC1 and apoptosis signal–regulating kinase 1 (ASK1) was also observed. RAC1 activation contributes to oxidative stress and inflammatory signaling in hepatocytes (105), while ASK1 is a stress-responsive kinase implicated in steatohepatitis progression (106,107). Inhibition of phosphorylated CYP2E1 further supports attenuation of oxidative stress burden, as CYP2E1 is a major source of reactive oxygen species in steatotic liver and drives lipid peroxidation and mitochondrial dysfunction (108,109). Additionally, suppression of pro-apoptotic mediators BAX and caspase-7 (CASP7) indicates attenuation of intrinsic apoptotic signaling, a hallmark of progressive NAFLD and fibrogenic remodeling (110,111).

To mechanistically contextualize the imaging observations, application of the Proteas BioTAS platform to plasma samples from the same STOP cohort identified significant inhibition of the NAFLD pathway at both total protein (DAP z-score = −1.37) and phosphorylation-dependent levels (DApP z-score = −1.61) (**Figure 8; Supplementary Table 11**). Pathway interrogation revealed convergent modulation of TNFα, insulin resistance, peroxisome proliferator–activated receptor (PPAR), adipocytokine, oxidative stress, and apoptotic signaling cascades—axes centrally implicated in progression from steatosis to steatohepatitis and fibrosis (95–98,110–112).

**Figure 8.**
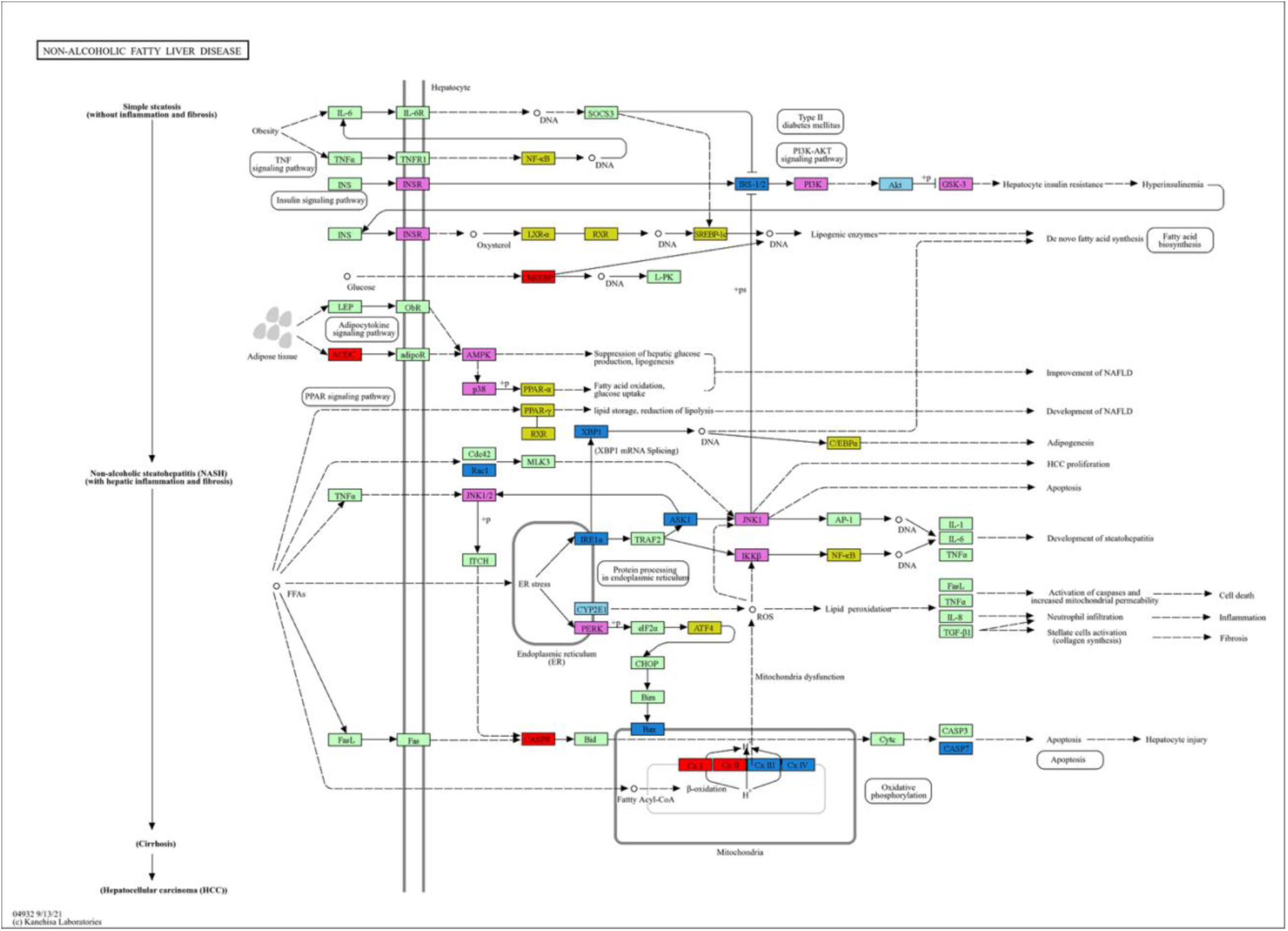
Non-alcoholic Fatty Liver Disease pathway mapping *(DEP z-score=-1.37; DApP z-score=-1.61)* was performed by overlaying differentially expressed proteins, predicted transcription factors, and predicted kinases onto KEGG pathways and visualized by category: upregulated proteins (red), downregulated proteins (blue), upregulated phosphoproteins (orange), downregulated phosphoproteins (sky blue), predicted transcription factors (yellow), and predicted kinases (purple). Statistical significance was determined using chi-squared testing with Benjamini–Hochberg false discovery rate (FDR) correction (q = 2.16 × 10^-8^). Predicted transcription factors and kinases were identified through overrepresentation enrichment analysis of the differentially expressed proteins.

The importance of these circulating pathway-level findings is reinforced by recent translational work demonstrating that semaglutide directly modulates metabolic, inflammatory, and fibrotic pathways in metabolic dysfunction–associated steatohepatitis (MASH). In a comprehensive multi-omic and histopathologic analysis, Jara et al. showed that semaglutide therapy resulted in coordinated downregulation of inflammatory and fibrogenic signaling networks, improvement in steatohepatitis activity, and attenuation of pathways linked to insulin resistance and oxidative stress (112). The convergence between tissue-level pathway modulation reported in that study and the plasma proteomic/phosphoproteomic signatures identified here substantiates the biological validity of the BioTAS-derived circulating markers as reflective of intrahepatic disease remodeling.

These findings are further supported by prior application of the methodological precepts of the BioTAS platform in the randomized WELCOME trial of patients with NAFLD treated with pharmacologic grade high-dose (4g per day) marine omega-3 fatty acids (26), in which plasma proteomic profiling demonstrated coordinated modulation of inflammatory, oxidative stress, and thrombosis-associated pathways paralleling improvements in hepatic steatosis. Together, these data underscore the translational robustness of plasma phosphoproteomics for mechanistic interrogation of NAFLD and its cardiometabolic sequelae.

Importantly, NAFLD is increasingly recognized as a multisystem disease strongly associated with atherosclerotic cardiovascular disease (113). The coordinated suppression of hepatic inflammatory, oxidative, and apoptotic signaling observed here occurred in parallel with reductions in epicardial and hepatic fat (6,8), inhibition of lipid-driven inflammatory cascades (80–94), and attenuation of AGE–RAGE signaling (52–57,60).

### Coordinated Inhibition of the Diabetic Cardiomyopathy (DCM) Signaling Network

Building upon the integrated suppression of AGE–RAGE, insulin resistance, lipid–atherosclerosis, and NAFLD pathways described above (52–57,60,69–79,80–94,95–113), semaglutide treatment in the STOP cohort was associated with coordinated inhibition of the diabetic cardiomyopathy (DCM) signaling network. GLP-1 receptor agonists (GLP-1RAs), including semaglutide and liraglutide, have emerged as therapeutic agents capable of mitigating key features of DCM—a myocardial disorder characterized by increased ventricular stiffness, impaired diastolic relaxation, and progressive systolic dysfunction occurring independently of overt coronary artery disease or hypertension. Experimental and clinical studies demonstrate that GLP-1RAs improve myocardial energetics, reduce oxidative stress, and attenuate inflammatory and fibrotic remodeling in the diabetic myocardium (114,115).

Consistent with this body of evidence, our proteomic and phosphoproteomic analysis demonstrated significant inhibition of the composite DCM pathway following semaglutide treatment compared with placebo (DAP z-score = −1.32; DApP z-score = −2.15; **Figure 9, Supplementary Table 12**). This pathway reflects coordinated dysregulation of multiple interdependent signaling axes central to DCM pathogenesis, including AGE–RAGE signaling (52–57,60), mitochondrial tricarboxylic acid (TCA) cycle flux, myocardial glucose oxidation, TGF-β–mediated fibrotic signaling, insulin signaling (69–74), and activation of the renin–angiotensin–aldosterone system (RAAS). Persistent activation of these networks promotes metabolic inflexibility, mitochondrial dysfunction, extracellular matrix deposition, and chronic low-grade inflammation within the diabetic heart (114,115).

**Figure 9.**
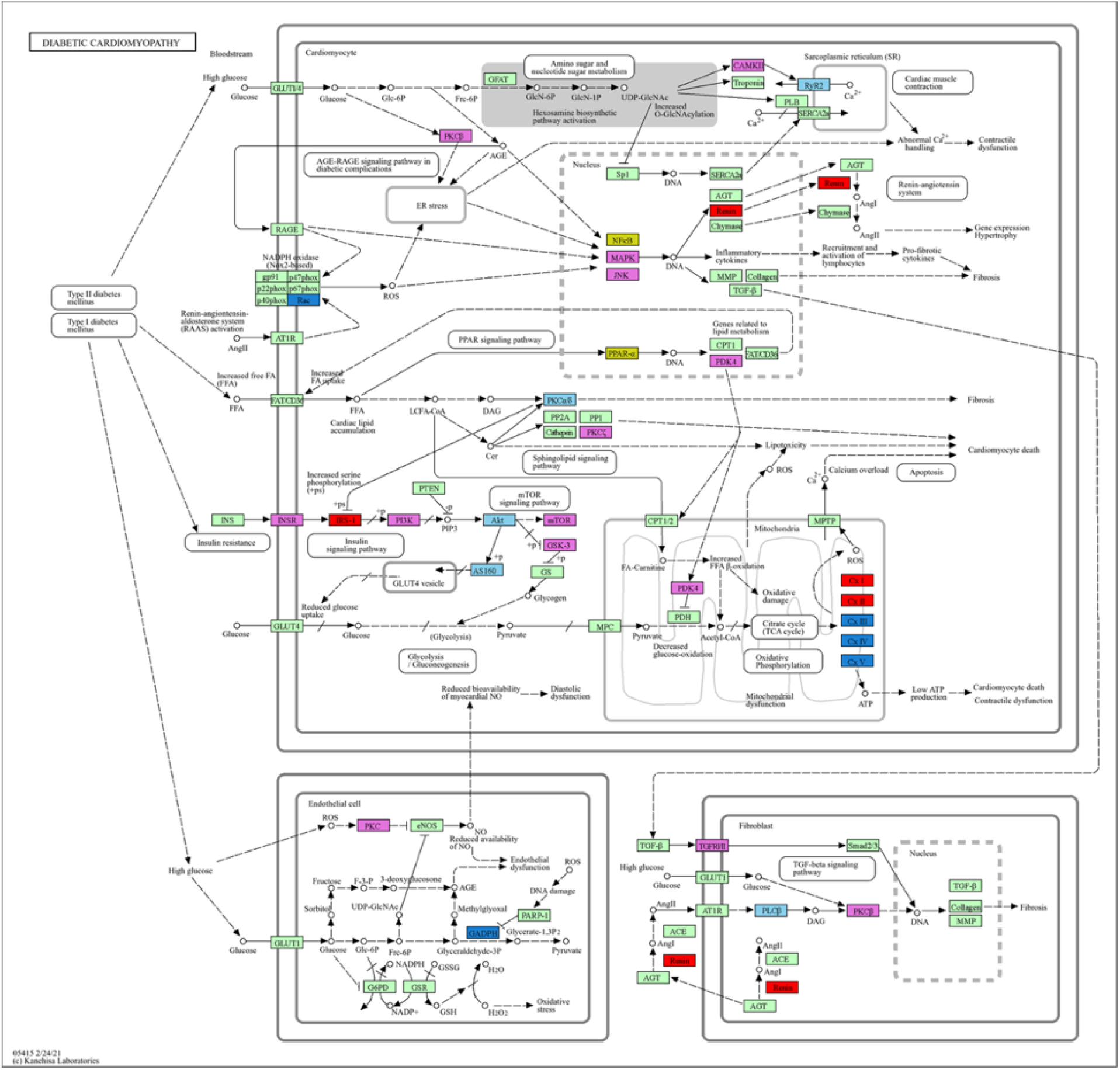
Diabetic Cardiomyopathy pathway mapping *(DEP z-score=-1.32; DApP z-score=-2.15)* was performed by overlaying differentially expressed proteins, predicted transcription factors, and predicted kinases onto KEGG pathways and visualized by category: upregulated proteins (red), downregulated proteins (blue), downregulated phosphoproteins (sky blue), predicted transcription factors (yellow), and predicted kinases (purple). Statistical significance was determined using chi-squared testing with Benjamini–Hochberg false discovery rate (FDR) correction q = 7.27 × 10^-3^). Predicted transcription factors and kinases were identified through overrepresentation enrichment analysis of the differentially expressed proteins.

AGE–RAGE signaling represents a critical upstream driver of oxidative stress, inflammatory cytokine production, and profibrotic gene expression in DCM, linking hyperglycemia-induced protein glycation to myocardial stiffening and contractile dysfunction (52–54,60,116,117). Experimental attenuation of AGE accumulation or RAGE silencing improves myocardial compliance, reduces reactive oxygen species (ROS) generation, and limits extracellular matrix remodeling (116,117). The observed suppression of the integrated DCM pathway in our study is therefore mechanistically concordant with inhibition of AGE–RAGE–dependent downstream signaling and aligns with the coordinated anti-inflammatory and metabolic remodeling observed across hepatic, vascular, and adipose compartments.

Importantly, GLP-1 receptor activation has also been shown to favorably modulate myocardial substrate utilization, shifting metabolism toward improved mitochondrial efficiency and enhanced fatty acid oxidation, while attenuating glucotoxic and lipotoxic stress (115,118). In parallel with the insulin-sensitizing phospho-signatures described above (69–74), these myocardial effects support restoration of metabolic flexibility—a key defect in DCM.

An additional observation from the BioTAS analysis was a reduction in circulating glyceraldehyde-3-phosphate dehydrogenase (GAPDH) abundance in the semaglutide-treated group. Beyond its canonical glycolytic role, GAPDH functions as a redox sensor and mediator of apoptotic and nuclear transcriptional signaling (119). Decreased GAPDH abundance has been reported during fasting and fasting-mimetic states, including GLP-1RA–associated metabolic reprogramming (69–71), and may reflect a shift away from excessive glycolytic flux toward enhanced oxidative metabolism also observed in carcinogenesis (120).

Notably, reduction of GAPDH in the present study occurred in concert with inhibition of mTORC1 signaling. mTORC1 activation has been implicated in pathological cardiac hypertrophy, metabolic dysregulation, and increased glycolytic protein synthesis (121). Prior quantitative proteomic analyses by the authors using the QuaNCAT LC-MS method demonstrated that hypertrophic stimulation in primary cardiomyocytes induces *de novo* GAPDH synthesis in an mTORC1-dependent manner, verified by mRNA and immunoblot analyses (122). The reciprocal reduction of GAPDH and mTORC1 signaling observed here is therefore consistent with attenuation of hypertrophic and glycolysis-driven metabolic remodeling, diminished ROS generation, and protection against glucotoxic myocardial injury.

To our knowledge, these findings represent the first demonstration of coordinated suppression of the DCM signaling network at the systemic plasma proteome and phosphoproteome level in response to GLP-1RA therapy. This systems-level molecular signature extends beyond glycemic control and complements emerging clinical data demonstrating reductions in heart failure–related outcomes among patients treated with GLP-1RAs (62,63,115).

#### Discussion and future perspectives

Plasma represents a uniquely informative and minimally invasive window into human health and disease, integrating molecular signals derived from vascular, hepatic, adipose, myocardial, and immune compartments (123–129). Large-scale proteogenomic investigations have demonstrated that circulating proteins reflect both inherited regulation and active pathophysiologic processes across organ systems (125, 126, 129, 130). However, effective monitoring of disease initiation, progression, and therapeutic response requires unbiased interrogation of protein abundance together with post-translational modification status—particularly phosphorylation, which defines active signaling states and therapeutic target engagement (131–133). This objective is technically demanding. The plasma proteome is extraordinarily complex: the ∼20 most abundant proteins account for >98% of total protein mass, while native protein concentrations span more than 10 orders of magnitude (123,124). In addition, pre-analytical variables—including collection, processing delays, hemolysis, and storage conditions—can significantly alter protein abundance and phospho-status, introducing bias (134–136). These constraints necessitate high-dynamic-range, phosphorylation-aware analytical platforms.

Affinity-based proteomic technologies, although enabling multiplex throughput, are intrinsically constrained by predefined reagent panels, epitope dependence, and incomplete proteome representation (125, 126, 127, 134–140). They do not resolve phosphorylation-dependent functional states central to cardiometabolic signaling biology (131–139), and protein-altering variants or matrix effects may influence measurements independently of true biological abundance (125,126, 128, 129). Recent technical benchmarking studies highlight substantial differences in proteome depth, quantitative performance, and post-translational modification coverage across plasma proteomics platforms, underscoring limitations of predefined panel approaches relative to unbiased LC–MS (140).

Unbiased multidimensional LC–MS strategies overcome many of these constraints. Discovery plasma proteomics has consistently demonstrated that substantial fractions of the circulating and extracellular vesicle proteome are not captured by affinity-based assays (30, 124, 127, 135, 140, 141). In a seminal JCI Insight study, Garay-Baquero et al. reported an unprecedented 11-orders-of-magnitude absolute linear dynamic range in human plasma—the largest reported to date—enabling detection of extremely low-abundance, disease-relevant regulatory proteins enriched in extracellular vesicles and undetectable by antibody- or aptamer-based platforms (30). This study applied earlier multidimensional LC–MS methodological principles (20,21,23) that underpin the technological basis of the patented BioTAS platform.

In the STOP randomized trial, BioTAS extended these discovery principles into a clinical setting by interrogating plasma samples matched to serial coronary computed tomography angiography (CCTA) and cardiac CT–based metabolic imaging assessments. In semaglutide-treated patients, phosphoproteomic profiling demonstrated coordinated suppression of AGE–RAGE, insulin resistance, lipid–atherosclerosis, NAFLD, and diabetic cardiomyopathy signaling networks. These molecular changes occurred alongside significant reductions in blood glucose and HbA1c levels relative to placebo, confirming improved glycemic control. Imaging analyses further demonstrated reductions in epicardial adipose tissue volume, attenuation of liver fat content, and favorable coronary plaque characteristics by CCTA (6–8).

The circulating proteomic and phosphoproteomic remodeling captured by BioTAS closely recapitulated established experimental and clinical evidence that GLP-1 receptor agonists attenuate oxidative stress, inflammatory activation, lipotoxicity, and fibrotic remodeling (52–57,60,62,63,69–74,80–94,95–118). The convergence of three orthogonal modalities—biochemical improvement (glucose and HbA1c), structural and metabolic remodeling by advanced imaging, and phosphorylation -resolved pathway suppression in plasma—provides compelling cross-validation of the molecular findings. This integrated molecular–biochemical–imaging concordance substantiates the analytical validity, biological specificity, and translational robustness of the BioTAS platform.

As a perspective, comprehensive BioTAS profiling of the entire STOP cohort—and extension to larger observational studies and randomized controlled trials evaluating next-generation GLP-1 receptor agonists, including oral formulations—offers a transformative opportunity for precision cardiometabolic therapeutics. Beyond discovery, translation into clinical practice requires analytically robust quantitative validation. Targeted mass spectrometry assays, which rely on high-throughput benchtop triple quadrupole designs, are increasingly recognized within clinical laboratory frameworks as precise, reproducible, and scalable modalities for laboratory-developed test implementation, and evolving regulatory landscapes provide structured pathways for validation of multiplex LC–MS–based diagnostics (142–149).

Accordingly, mechanistically selected BioTAS-derived proteins and phosphoproteins can be incorporated into a gold-standard targeted LC–MS companion diagnostic panel. Integration of optimized sample procurement protocols with nanotechnology-enhanced protein and phosphoprotein isolation, separation, and enrichment modalities intrinsic to the BioTAS workflow would enable highly sensitive and phosphorylation-aware quantification across a broad dynamic range. Such a platform would support precision titration of GLP-1RA monotherapy, rational evaluation of pharmacologic synergists, and integration with lifestyle adjuvants—including nutrition and structured physical activity (150,151).

The interpretive complexity of high-dimensional phosphoproteomic datasets further benefits from AI-assisted frameworks capable of integrating evolving biological knowledge with kinase–substrate network modeling (152). These approaches enhance mechanistic interpretability and accelerate translation from systems-level discovery to individualized clinical decision support.

Ultimately, implementation of a validated, targeted LC–MS–based GLP-1RA companion diagnostic panel grounded in BioTAS discovery principles would represent a major advance toward personalized cardiometabolic medicine. By enabling dynamic monitoring of pathway engagement, safety signals, and mechanistic efficacy across diverse therapeutic regimens—including emerging oral GLP-1RAs and combination strategies—this integrated molecular platform addresses a critical unmet need in the pharmacologic industry for precision-guided, optimally safe, and efficacious cardiometabolic intervention.

As a proof-of-concept investigation, the present study was intentionally focused on pathway-level results that could be directly aligned with imaging findings obtained from matched patients randomized to semaglutide versus placebo. This design allowed independent validation of the clinical accuracy and biological specificity of the BioTAS platform by demonstrating concordance between circulating proteomic/phosphoproteomic remodeling and structural phenotypes assessed by advanced imaging.

Beyond the pathways explicitly presented in conjunction with imaging endpoints, multiple additional signaling networks were quantified with strong statistical significance. These networks were functionally linked to coordinated inhibition of the AGE–RAGE and insulin signaling cascades—two central mechanistic axes identified in this analysis. Given the well-established role of AGE–RAGE activation and insulin resistance in systemic inflammatory–metabolic dysregulation, suppression of these upstream drivers has broader implications extending beyond coronary and hepatic remodeling.

Specifically, the pathway signatures observed in this cohort intersect with biological processes implicated in diabetic sarcopenia, osteopenia, retinopathy, aging-related tissue degeneration, and neurocognitive decline. Although these phenotypes were not primary endpoints of the STOP trial, the statistically robust modulation of their underlying signaling pathways supports a systems-level disease-modifying effect of GLP-1 receptor agonist therapy. These findings warrant further investigation in larger cohorts and dedicated clinical studies to determine whether the molecular signatures identified by BioTAS translate into measurable improvements in musculoskeletal, retinal, and neurocognitive outcomes.

## Acknowledgements

The authors thank Novo Nordisk for critical manuscript review. The authors declare that the BioTAS platform technology and derivative cardiometabolic signatures applied for this study is associated with Patent Pending No. PCT/US2021/063407.

## Conflict of interest

A.M. is co-Founder and CSO of Proteas Health, Inc. R.N. is CEO of Proteas Health, Inc. and S.D.G. is Founder, President and CTO of Proteas Health, Inc.

## Funding Information

This work was supported by an NIH NHBLI SBIR Phase 1 grant (no.13525094) and Proteas Health, Inc.

