## Supplementary Methods for "Novel Plasma Proteomics and Phosphoproteomics Platform Captures Pleiotropic Cardiometabolic Spectrum Effects of Semaglutide in Patients with T2D and Atherosclerosis: A Companion Diagnostic Pilot Study from the STOP (Semaglutide Treatment On coronary atherosclerosis Progression) Randomized Trial"

Antigoni Manousopoulou, et al.

**SUPPLEMENTARY METHODS**

*Plasma procurement:* The plasma samples were obtained from the clinical cohort examined in the STOP randomized trial (7). Specifically, 8 patients in the active group at baseline and 52 weeks post treatment and 8 patients in the placebo group at baseline and 52 weeks post treatment were subjected to proteomic analysis. The clinical characteristics of the cohort at baseline are presented in Tables 1-3. A volume of 100μL of whole plasma from each specimen was aliquoted and vortex mixed for 30 sec with 400μL of Proteas’ proprietary liquid fixative solution before analysis and also served as the mobile phase for the fractionation of its protein content. This chaotropic liquid fixative also include viscosity modifiers, detergents and organic solvents that affords maximum hydrodynamic radii for the plasma proteins/phosphoproteins, as required for their optimum partitioning and tailored to work with acrylic glass monolithic micro-column described below. The liquid fixative also allows for the neutralization of plasma endogenous proteases, phosphatases, phosphotransferases and the solubilization of membranous proteins and phosphoproteins that constitute lipid microvessicles, such as exosomes. An additional attribute is the sterilization of all possible blood-borne pathogens such as Hep A, B, C; HIV, COVID-19 and TB as verified with microbiological testing. The proteome composition of the liquid fixative treated plasma samples have been verified to remain stable for a minimum of 72 hours at temperatures between 18-25^0^C, up to 3-months between 4 – 8^0^C, and over 4 years at -20^0^C based on stability studies conducted using the same proteomics platform and methods, as described in this study.

*Proteomic sample preparation:* A total volume of 80 μL from each liquid fixative plasma extract was subjected automated serial fractionation for its protein and native peptide content using a variant of the open tubular chromatography array lab-on-chip originally reported by the authors (39) and most recently subject matter of Proteas international patent (17). This lab-on-chip is an acrylic glass monolithic micro-column with 32 parallel channels each at 3.8 cm length x 7.5 mm width x 40 mm depth generated with photolithography and deep plasma etching. An isocratic flow rate of 10μL/min with column temperature maintained at 50ºC. The injection volume for each iteration was 20 μL for a total of 4 iterations to achieve~~d~~ the desired serial concentration enrichment. Furthermore, the etched morphology of the bottom surface is constituted of an array of macropores and mesopores that enabled the effective partitioning of the plasma proteins, provided that the samples injection volume remains less than 20μL for each iterative run. The liquid fixative composition facilitated the fractionation process given its high chaotropic properties and tailored viscosity properties to enable sufficient interaction time between the protein constituents and the inner domain of lab-on-chip under the defined flow rate, injection volume and column temperature. The partitioning efficiency was evaluated with standardized protein mix and reference liquid fixative preserved plasma extracts. The lab-on-chip was enclosed in device holder (Micronit, device holder no 4515) for its interfacing with bioinert Nexera HPLC system (Shimadzu, Canby, Oregon). Poly-imide inner coated fused Silica capillary tubing with 150 mm ID and and 360 mm OD coated, and related connectors, to the chip holder were used (Upchurch Scientific). Eluents were monitored at 218nm and 280nm and collected at predefined fractionation times at 4ºC using the Spider 10-port configuration as reported by the authors (40), and repurposed per the Proteas patent (17). The eluents were collected in mini-dialysis cups (Thermo Fisher Scientific) and were eventually exchanged with 0.5 M TEAB and 0.1% SDS at 4ºC using a customized continuous high flux buffer exchange system, measured for its protein content and subjected to reduction, alkylation and LysC/Trypsin proteolysis at a ratio of 1:30. The resulting proteotypic peptides/phosphopeptides were labeled with the TMTpro isobaric stable isotope reagents, as specified by the manufacturer (Thermo Pierce, Rockford, IL).

*Phosphopeptide isolation and enrichment*: The TMTpro labeled peptide/phosphopeptide samples was subjected to step-gradient chromatography with an open tubular Lab Chip surface functionalized with TiO2/ZrO2 chemistry as reported by the authors (39). A total volume of 40μL from each sample was interactively injected to TiO2/ZrO2 LabChip in 10μL volume increments. The initial mobile composition was 0.1% aqueous formic acid (pH 3) at a flow rate of 10μL/min for 15min to allow binding of its phosphopeptide content while allowing the non-phosphorylated peptides to elute and collected for follow-up LC-MS analysis. To facilitate this process, the column temperature was maintained at 50 ºC. The bound phosphopeptides were eluted with the introduction of the alkaline mobile phase constituted of 0.1 M NH_4_OH (pH 11) at a flow rate 10μL/min for 30min and column temperature maintained at 50 ºC. The eluted non-phosphorylated peptide samples were desalted with the HLB cartridges and were lyophilized down to dryness. The phosphopeptide enriched fractions were lyophilized down to dryness. All samples were reconstituted in MB and 0.2-micron particle filtrated before analysis.

*Mass Spectrometry Analysis:* Samples reconstituted in LC buffer A (0.1% formic acid in water), randomized, and then injected onto an EASY-nLC 1200 ultra-high-performance liquid chromatography coupled to a Exploris 480 quadrupole-Orbitrap mass spectrometer (Thermo Fisher Scientific) with internal calibration and retrofitted with nano-spray ionization source. To exclude +1 charged species, the Field Asymmetric Ion Mobility Source (FAIMS) was used at -40 and -70CV. Peptides were separated by a custom reverse phase analytical column integrated with nanospray emitter (Mixed mode C4 and C18, 1.6 µm particles, 100 Å pore, 50 µm ID × 25 cm L). Flow rate was set to 400 nL/min at a gradient from 3% LC buffer B (0.1% formic acid, 80% acetonitrile) to 38% LC buffer B in 110 min, followed by a 10-min washing step to 85% LC buffer B. The maximum pressure was set to 1,000 bar, and column temperature was maintained at 60°C. Peptides separated by the column were ionized at 1.5 kV in positive ion mode. MS1 survey scans were acquired at the resolution of 240,000 from 350 to 1,800 m/z, with a maximum injection time of 100 ms and AGC target of 1e6. MS/MS fragmentation of the 10 most abundant ions were analyzed at a resolution of 60,000, AGC target 5e4, maximum injection time 100 ms, and normalized collision energy of 32. Dynamic exclusion was set to 30 sec, and ions with charge +1, +4 and >+4 were excluded. Figure 1 illustrates an overview of the proteomic platform applied for this pilot study.

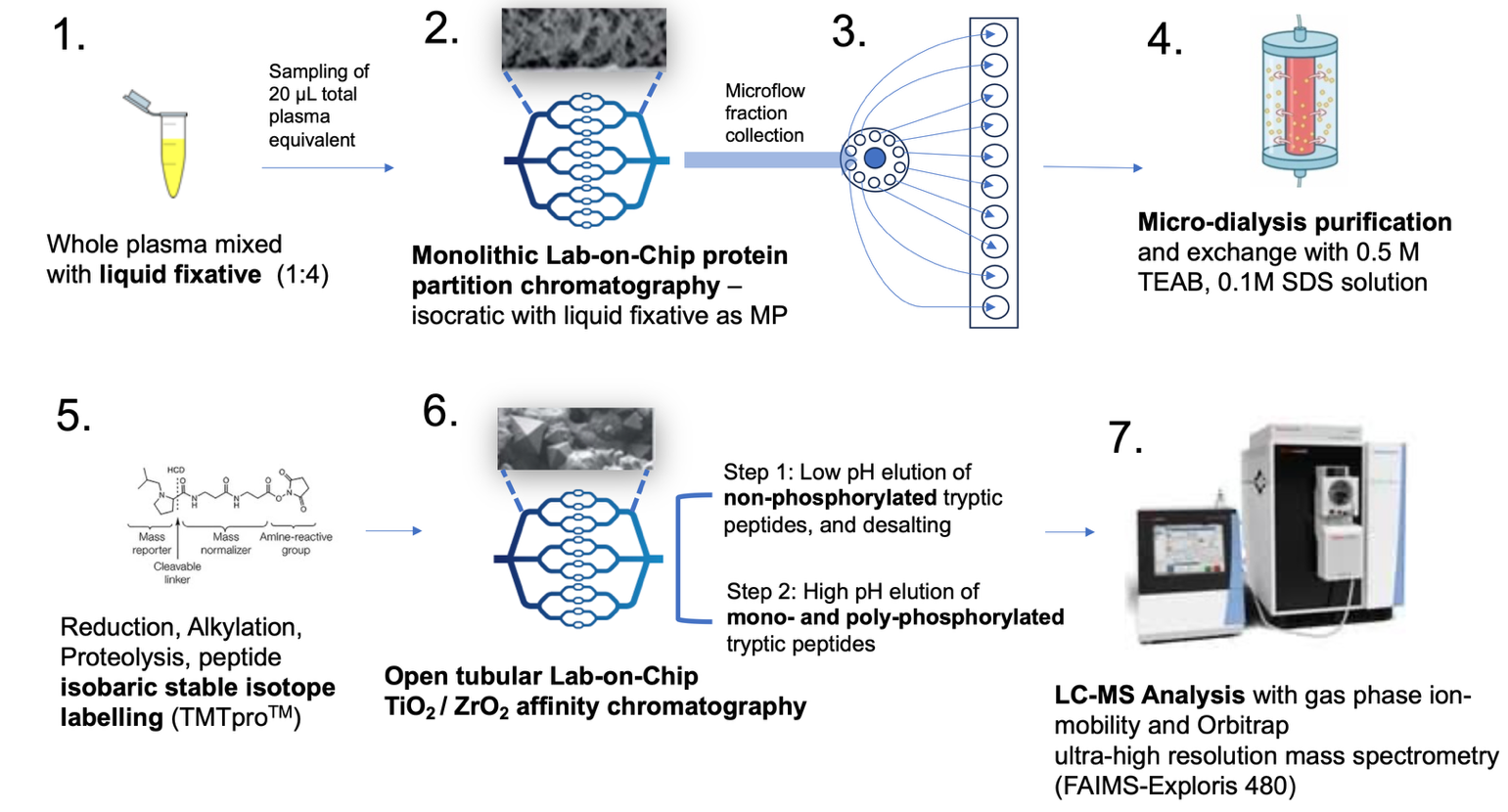

**Figure 1.** The basic steps of the **Bio**fluid **T**otal **A**nalytic **S**ystem (BioTAS) proteomics platform applied to study.
