## Supplementary Figures for "Novel Plasma Proteomics and Phosphoproteomics Platform Captures Pleiotropic Cardiometabolic Spectrum Effects of Semaglutide in Patients with T2D and Atherosclerosis: A Companion Diagnostic Pilot Study from the STOP (Semaglutide Treatment On coronary atherosclerosis Progression) Randomized Trial"

Antigoni Manousopoulou, et al.

**Supplementary Figures**

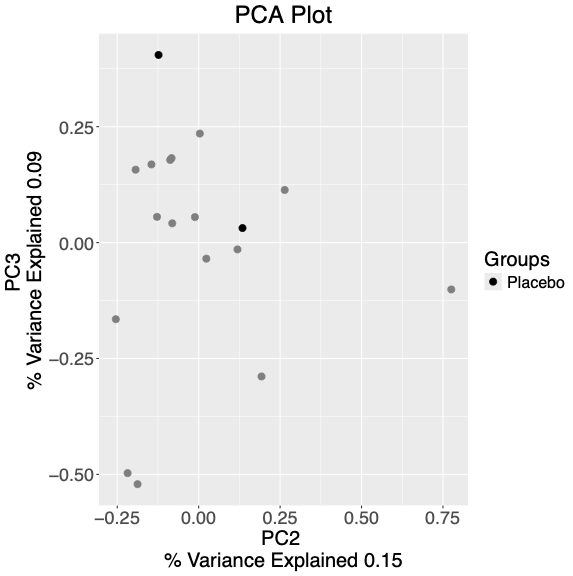
Protein and phosphorylated protein abundance values were log2 transformed and subjected to principal component analysis.  Principal components were ranked by total variance explained and plotted.  The Variance explained by each principal component is shown in the axes.  Coloration indicates groups for samples with purple for Semaglutide, green for Baseline, and black for placebo.

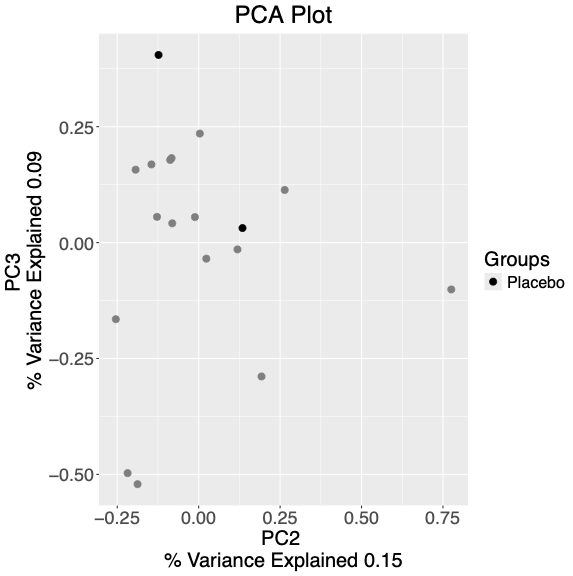

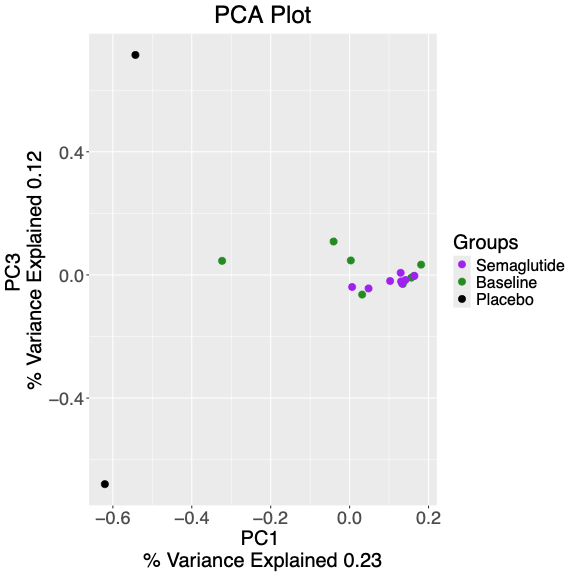

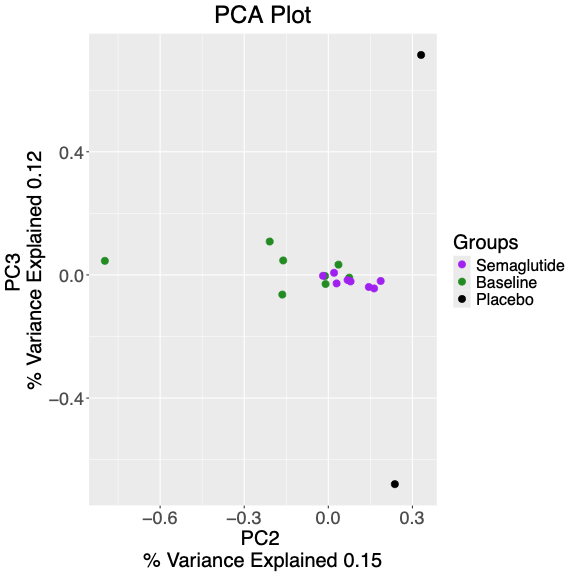

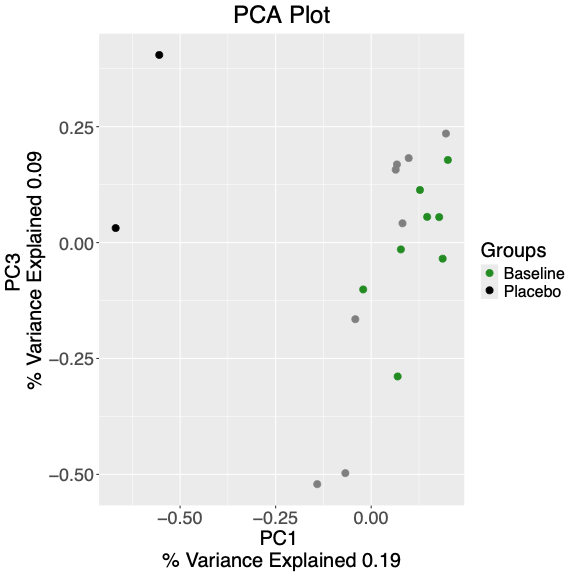
