## Supplementary Tables 1-4 for "Novel Plasma Proteomics and Phosphoproteomics Platform Captures Pleiotropic Cardiometabolic Spectrum Effects of Semaglutide in Patients with T2D and Atherosclerosis: A Companion Diagnostic Pilot Study from the STOP (Semaglutide Treatment On coronary atherosclerosis Progression) Randomized Trial"

**Supplementary Table 1**. Demographics of cohort at randomization

| Demographics | Semaglutide | Placebo | p-value |
|---|---|---|---|
| | n=8 | n=8 | |
| Age (years) | 47.3 ± 3.9 | 49.0 ± 5.2 | 0.81 |
| Body Mass Index ($kg/m^2$) | 31.1 ± 4.7 | 32.3 ± 4.0 | 0.43 |
| Male (%) | 8 (100) | 8 (100) | 1 |
| Hispanic/Latino (%) | 5 (60) | 5 (60) | 1 |
| Race | | | 1 |
| White (%) | 7 (87.5) | 7 (87.5) | |
| Asian (%) | 1 (12.5) | 1 (12.5) | |
| Black or African American (%) | 0 (0) | 0 (0) | |
| Other (%) | 0 (0) | 0 (0) | |
| Follow-up time (years) | 1.1 ± 0.5 | 1.1 ± 0.5 | 0.848 |

**Supplementary Table 2**. Risk Factors of cohort at randomization

| Risk Factors | Semaglutide | Placebo | p-value |
|---|---|---|---|
| | n=8 | n=8 | |
| Hypertension (%) | 5 (62.5) | 5 (62.5) | 1 |
| Hyperlipidemia (%) | 7 (87.5) | 7 (87.5) | 1 |
| Family History of CAD (%) | 2 (25) | 2 (25) | 1 |
| Current Smoker (%) | 1 (12.5) | 1 (12.5) | 1 |
| Past Smoker (%) | 2 (25) | 2 (25) | 1 |

**Supplementary Table 3**. Laboratory values of cohort at randomization

| Laboratory Values | Semaglutide | Placebo | p-value |
|---|---|---|---|
| | n=8 | n=8 | |
| Mean Plasma Glucose (mg/dL) | 198.0 ± 55.0 | 205.0 ± 59.0 | 0.57 |
| Hemoglobin A1C (%) | 8.5 ± 1.5 | 8.8 ± 2.0 | 0.63 |
| Triglycerides (mg/dL) | 145.0 ± 65.0 | 138.0 ± 60.2 | 0.35 |
| HDL-C (mg/dL) | 39.4 ± 10.0 | 39.0 ± 9.9 | 0.85 |
| LDL-C (mg/dL) | 74.2 ± 31.5 | 76.5 ± 30.5 | 0.38 |
| hsCRP (mg/L) | 1.5 (0.7, 5.0) | 2.2 (0.8, 4.5) | 0.25 |
